# Spatially Confined Phase Separation Promotes Local Tubulin Availability to Stabilize Ciliary Microtubule Growth

**DOI:** 10.64898/2026.09.14.751438

**Authors:** Kaiming Xu, Zhiwen Zhu, Yongping Chai, Wei Li, Guangshuo Ou

## Abstract

How microtubule (MT) growth is locally sustained within cilia remains a central unanswered question in ciliogenesis. Here we identify RAT-1 as a regulator of ciliary tubulin homeostasis and MT growth. Genetic screens in *Caenorhabditis elegans* show that loss of RAT-1 suppresses the toxic effects of a mutant α-tubulin and restores ciliary structure, while RAT-1 deficiency reduces tubulin turnover at the ciliary tip, indicating its physiological role in tuning ciliary MT dynamics. Mechanistically, RAT-1 forms intra-ciliary condensates selectively at the distal region through an auto-inhibitory α-helix and phosphorylation by the distally localized ciliary kinase. RAT-1 enriches tubulin within condensates in vivo/vitro, and cooperates with end-binding proteins to facilitate sustained MT growth in vitro. Notably, heterologous expression of worm RAT-1 reveals conserved condensate behavior within mammalian cilia. Our findings reveal a kinase-regulated mechanism controlling RAT-1 condensation within cilia and establish RAT-1 as a regulator of ciliary MT dynamics.

## Introduction

Cilia are microtubule (MT)-based organelles protruding from the surface of most eukaryotic cells, performing essential functions in signal transduction, environmental perception, and motility (Anvarian et al., 2019; Hilgendorf et al., 2024; Klena and Pigino, 2022; Lacey and Pigino, 2025; Ou and Scholey, 2022). Structurally, cilia comprise 9 MT doublets, each composed of a complete A-tubule with 13 protofilaments and an incomplete B-tubule containing 10 protofilaments (Chen and Ou, 2025; Lacey and Pigino, 2025; Ma et al., 2019; Walton et al., 2023). Cilia are tiny, crowded compartments, yet the MTs must grow precisely and continuously at the ciliary tips (Brouhard and Rice, 2018; Gudimchuk and McIntosh, 2021; Saunders et al., 2025). How tubulins—the fundamental building blocks of MTs—are delivered and locally tuned within this confined space has remained a long-standing mystery (Akhmanova and Kapitein, 2022; Janke and Magiera, 2020).

Ciliary tubulins are transported as cargoes to the ciliary tip via intraflagellar transport (IFT) (Bhogaraju et al., 2013; Craft et al., 2015; Kubo et al., 2016; Lacey and Pigino, 2025; Ou and Scholey, 2022; Taschner et al., 2016). Once inside the cilia, tubulins are unloaded from the IFT complexes and then incorporated into ciliary MTs (Jiang et al., 2022; Maurya et al., 2019). While IFT ensures efficient delivery, how cilia maintain a readily accessible pool of tubulins at the distal tip to support continuous MT elongation, and how this pool is stored or mobilized dynamically, remains largely unknown. Elucidating these processes is of critical importance, as perturbations in tubulin supply or MT dynamics compromise ciliary integrity and contribute to a wide spectrum of ciliopathies (Dodd et al., 2024; McKenna et al., 2023; Mollica et al., 2025; Reiter and Leroux, 2017).

Recent studies have revealed that intracellular phase separation can generate dynamic, membrane-less condensates that locally concentrate proteins and modulate biochemical reactions within spatially confined compartments (Chin Sang et al., 2025; Hadarovich et al., 2025; Qiu et al., 2024). Phase separation has emerged as a powerful organizer of cytoskeletal components, modulating polymerization dynamics and facilitating the rapid turnover of essential molecules. For example, BuGZ promotes spindle apparatus assembly by buffering tubulin through condensate formation (Jiang et al., 2015); SPD-5 functions as a scaffold to facilitate the assembly of pericentriolar material (PCM), a supramolecular membrane-less organelle (Woodruff et al., 2015). In the context of ciliogenesis, DynAPs (dynein axonemal particles), FGMs (fibrogranular materials), and centrosomal protein networks—including CAMSAP, WDR47, CEP164—can concentrate various ciliary proteins at the root of cilia to facilitate ciliogenesis (Chou et al., 2025; Huizar et al., 2018; Imasaki et al., 2022; Ren et al., 2022; Zhao et al., 2021). A recent study identified that phase separation of Spef1 is critical for central-pair MT functionality and motile ciliary beating (Ren et al., 2026). These findings underscore the indispensable role of phase separation during ciliogenesis. However, whether a similar mechanism operates within cilia to regulate tubulin availability, and how it might be spatially confined to the distal tip, remains unexplored. Moreover, little is known about how local kinase activity could regulate condensate formation to coordinate tubulin delivery and MT growth precisely (López-Palacios and Andersen, 2023).

Here, we identify RAT-1 (*Regulator of Alpha-Tubulin 1*), an unexplored ciliary protein, as a phase separation-mediated regulator of tubulin availability at the distal ciliary tip. RAT-1 is synthesized in the cytoplasm and transported along cilia via IFT but forms condensates selectively at the distal tip due to the presence of an internal inhibitory α-helix. Phosphorylation by the distally localized ciliary kinase DYF-5/MAK (Burghoorn et al., 2007) relieves this inhibition, thereby promoting local condensate formation. Functionally, RAT-1 concentrates tubulin within condensates and cooperates with end-binding (EB) family proteins to suppress catastrophe and enhance ciliary MT elongation. Together, these findings reveal a kinase-regulated, spatially confined phase separation mechanism that may couple tubulin supply to MT growth, uncovering a previously unrecognized principle of cytoskeletal regulation within cilia.

## Results

### RAT-1 is a tubulin-specific genetic suppressor required for ciliary integrity

To identify regulators of ciliary MT stability, we performed a forward genetic suppressor screen using the nematode *Caenorhabditis elegans* carrying TBA-5 (A19V), a ‘toxic’ and ‘gain-of-function’ mutation in ciliary α-tubulin (Fig. 1A) (Hao et al., 2011). The *tba-5 (A19V)* mutants lack ciliary distal segments at 15 ℃ and exhibit 100% dye-filling defective (Dyf) phenotype (Hao et al., 2011). From this background, we isolated suppressor strains that exhibited restored dye-filling capacity (Fig. 1A). Sibling subtraction method was performed to clone the suppressor mutation (Fig. S1A) (Joseph et al., 2018). From this screen, we isolated *rat-1 (F41E7.9),* also termed “Regulator of Alpha-Tubulin 1”, carrying a premature termination codon mutation (Q329*), which robustly rescued the Dyf phenotype of *tba-5 (A19V)* animals (Fig. 1A). This suppression was fully reversed by reintroduction of *rat-1* genomic DNA, or *gfp::rat-1* cDNA (Fig. 1B), confirming *rat-1* as the bona fide suppressor gene.

**Fig. 1.**
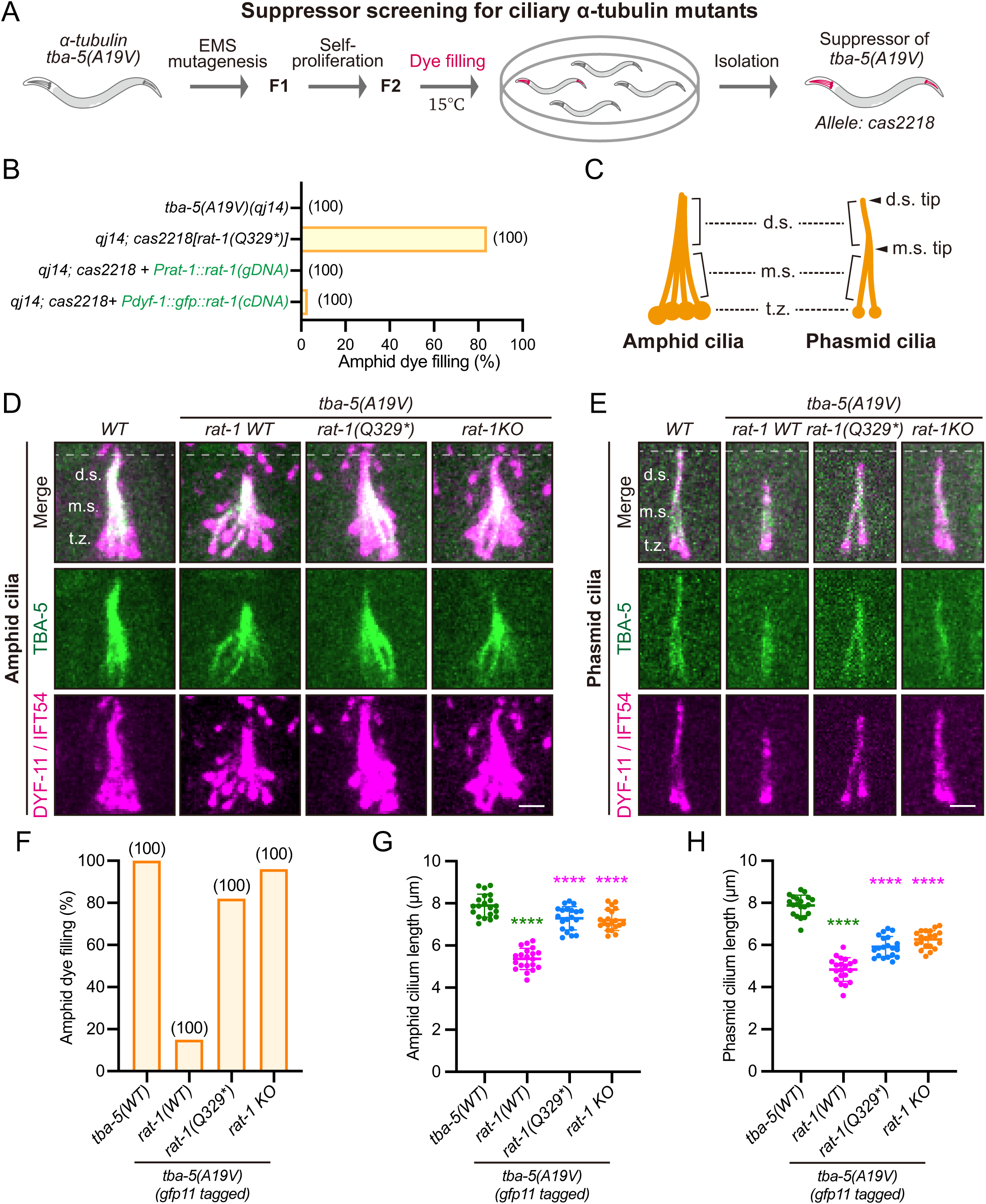
Loss-of-function *rat-1* rescues *tba-5 (A19V)*-induced ciliary microtubule defect. (A) Schematic of the forward genetic suppressor screen performed in the nematode *Caenorhabditis elegans* to identify suppressors of the ciliary defects caused by the α-tubulin mutant *tba-5 (A19V)*. Ethylmethanesulfonate (EMS) mutagenesis was performed to introduce genetic mutations into the genome of worms. Dye-filling assay was employed to evaluate the functionalities of cilia. (B) Percentage of amphid (containing sensory cilia in the head) dye-filling positive animals in each group. *cas2218* corresponds to the allele of the suppressor. The genome DNA (gDNA) sequence of *rat-1 (F41E7.9)*, or the cDNA sequence of *rat-1*, was transgenically overexpressed and indicated in green. (C) Schematic of *C. elegans* amphid and phasmid cilia. t.z., transition zone; m.s., middle segment; d.s., distal segment. m.s. tip and d.s. tip are indicated by black arrowheads. (D-E) Representative images of amphid (D) and phasmid (E) cilia in wild-type (WT) or mutant animals. *rat-1 KO* allele was created by genome editing (see also Fig. S1D). Cilia are marked with DYF-11(IFT54)::mScarlet. t.z., transition zone; m.s., middle segment; d.s., distal segment. Scale bar, 2 μm. (F) Percentage of amphid dye-filling positive animals in each group in (D). N = 100 animals per group. (G-H) Quantification of amphid (G) or phasmid (H) cilia length in each group. N = 20 phasmids. Shown as mean ± SD. One-way ANOVA with Dunnett’s tests were performed. **** *P* < 0.0001.

To examine the endogenous localization of RAT-1, we generated an N-terminal GFP knock-in allele at the *rat-1* locus using CRISPR-Cas9 genome editing (Figs. S1B-C). Wild-type GFP::RAT-1 exhibited robust ciliary localization, whereas GFP-tagged *rat-1 (Q329*)* animals displayed negligible fluorescence (Fig. S1C), confirming the suppressor allele is a loss-of-function mutation. We also generated an independent *rat-1* knockout allele (Fig. S1D). To visualize the endogenous localization of α-tubulin TBA-5 (A19V), we utilized a split-GFP-based functional tubulin labeling strategy (Fig. S1E) (Xu et al., 2024). Consistent with the rescue of *tba-5 (A19V)* Dyf phenotype, both *rat-1 (Q329*)* suppressor and the *rat-1* knockout restored amphid and phasmid ciliary length in split-GFP-tagged *tba-5 (A19V)* animals (Figs. 1C-H). Notably, loss of *rat-1* did not apparently impair the ciliary entry of TBA-5 (A19V) (Figs. 1D-E), consistent with the established role of IFT in tubulin delivery (Bhogaraju et al., 2013; Taschner et al., 2016).

Although *rat-1* loss suppressed *tba-5 (A19V)*-induced ciliary defects, *rat-1* knockout animals displayed shortened cilia, as revealed by fluorescence microscopy (Figs. S1F-G). While these animals retained normal dye-filling capacity (Fig. S1F), they exhibited partial resistance to ivermectin, an anti-nematode drug commonly used to assess ciliary dysfunction (Fig. S1H) (Campbell, 1993; Dent et al., 2000). Together, these findings establish RAT-1 as a tubulin-specific regulator required for normal ciliary structure and function.

### RAT-1 forms dynamic condensates at the distal region of cilia

AlphaFold3-predicted structural modeling of RAT-1 indicates the presence of an N-terminal intrinsically disordered region (N-IDR), a zinc finger-like domain (ZnFL) that lacks the canonical C2H2 or C2HC motif, a long internal α-helix (I-helix), and a C-terminal intrinsically disordered region (C-IDR) (Figs. 2A-B). To explore the intra-ciliary dynamics of RAT-1, we examined the intra-ciliary behaviors of RAT-1 in *gfp::rat-1* knock-in animals. In sensory cilia, GFP::RAT-1 localized as discrete condensates enriched in the distal region of cilia (Figs. 2C-D; Fig. S2A). Time-lapse imaging revealed that while the condensates were largely immobile, individual RAT-1 particles exhibited directed movement with velocities comparable to IFT and partial co-localization with IFT particles (Figs. S2B-C; Movie S1), suggesting RAT-1 is transported via IFT before accumulating distally. The highly disordered nature of RAT-1 (Fig. S2D) further supports the hypothesis that it undergoes phase separation within cilia. Indeed, fluorescence recovery after photobleaching (FRAP) showed rapid recovery of GFP::RAT-1 fluorescence within condensates (Figs. 2E-F; Fig. S2E), indicative of active molecular exchange and liquid-like behavior. Moreover, RAT-1 condensates occasionally exhibited fusion and fission-like behaviors within cilia (Figs. 2G-H; Movie S2), consistent with dynamic rearrangements of condensate structure in vivo. These findings suggest that RAT-1 may undergo liquid-liquid phase separation within cilia.

**Fig. 2.**
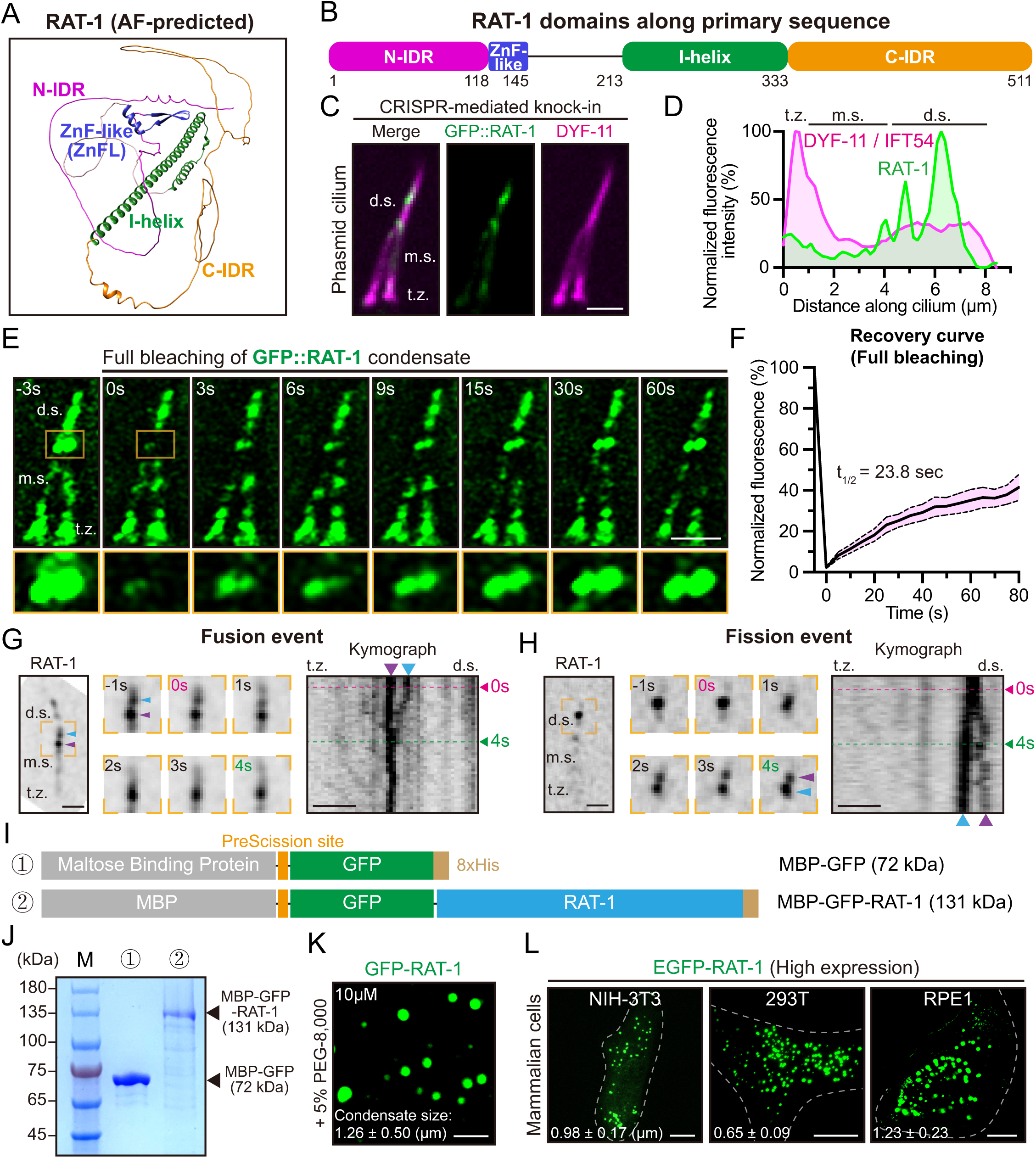
RAT-1 forms liquid-like condensates within the cilia and in vitro. (A) AlphaFold3 (AF)-predicted RAT-1 structure. The RAT-1 contains an N-terminal intrinsically disordered region (N-IDR), a zinc finger-like domain (ZnFL), an internal α-helix (I-helix), and a C-terminal intrinsically disordered region (C-IDR). (B) Distribution of RAT-1 domains along the primary sequence. (C) Representative image of phasmid cilium in *gfp::rat-1* knock-in animals. Cilia are marked with DYF-11(IFT54)::mScarlet. t.z., transition zone; m.s., middle segment; d.s., distal segment. Scale bar, 2 μm. (D) Distribution of RAT-1 and DYF-11 fluorescence along the cilium. The maximal flurescence was normalized to 100%. (E) Representative FRAP (fluorescence recovery after photobleaching) images of GFP::RAT-1 (knock-in) condensates within cilia. Scale bar, 2 μm. (F) Normalized fluorescence recovery of RAT-1 condensates within cilia after photobleaching. Shown as mean ± 95% CI (confidence interval). N = 10. (G) Representative fusion event of two GFP::RAT-1 (knock-in) condensates (marked in blue and purple, individually) within cilia. Kymograph was plotted along the cilium. Scale bar, 2 μm. See also Movie S2. (H) Representative fission event of RAT-1 condensates within cilia. Kymograph was plotted along the cilium. Scale bar, 2 μm. See also Movie S2. (I) Schematic of recombinant protein constructs for in vitro purification from bacteria. (J) SDS-PAGE analysis of purified proteins. MBP-GFP-RAT-1 and MBP-GFP (control) were loaded at equal total protein amounts as determined by Bradford assay. The gel shows the purity and molecular weight distribution of the purified proteins. M, marker. (K) Representative image showing GFP-RAT-1 (10 μM) forms liquid-like condensates in vitro. MBP (maltose binding protein) tag was removed by HRV 3C protease treatment, which cut the PreScission site specifically. Quantification of condensate size was shown as mean ± SD (N = 20 condensates). Scale bar, 5 μm. (L) Representative images showing overexpressed EGFP-RAT-1 forms liquid-like condensates in various mammalian cell lines (NIH-3T3, HEK293T and hTERT-RPE1). Quantification of condensate size was shown as mean ± SD (N = 20 condensates). Scale bar, 10 μm.

To test this hypothesis in vitro, we purified His-tagged MBP (maltose binding protein)-GFP-RAT-1 from bacteria (Figs. 2I-J) (see Methods). Consistent with in vivo observations, recombinant GFP-RAT-1 (10 μM) formed liquid-like condensates in vitro, in the presence of molecular crowding agent PEG (polyethylene glycol) (Fig. 2K). This condensate formation behavior depends on both protein and salt concentrations (Fig. S2F), and is strongly prohibited by phase separation inhibitor 1,6-hexanediol (Figs. S2G-H). After photobleaching, these RAT-1 condensates exhibit rapid fluorescence recovery in vitro (Figs. S2I-J). Additionally, transfection of EGFP-tagged RAT-1 into mammalian cell lines (NIH-3T3, HEK293T, and hTERT-RPE1) also resulted in robust condensate formation (Fig. 2L). Collectively, these results show that RAT-1 undergoes phase separation to form dynamic, liquid-like condensates, which are selectively enriched at the distal region of cilia.

### Domain architecture of RAT-1 underlies condensate formation

To investigate how RAT-1’s domains contribute to its function (Fig. 2B), we generated GFP-tagged truncation mutants and expressed them in *tba-5 (A19V); rat-1* knockout animals (Fig. S3A). Full-length RAT-1 rescued ciliary defects in *tba-5 (A19V)* single mutants, including reduced ciliary length and impaired dye-filling (Figs. 3A-B; Figs. S3B-C). In contrast, deletion of any major domain abolished this rescue, indicating loss of function (Figs. 3A-B; Figs. S3B-E). Fluorescence analysis showed that removal of the internal α-helix (I-helix) prevented ciliary entry and triggered premature condensate formation in dendrites, whereas deletion of the N-IDR, ZnFL, or C-IDR allowed ciliary entry but impaired condensate formation (Figs. 3A-D; Figs. S3D-E). These findings suggest that the N-IDR, ZnFL, and C-IDR are crucial for condensate formation, while the I-helix prevents premature condensation. Consistently, deletion of N-IDR, ZnFL or C-IDR eliminated condensate formation of RAT-1 in human RPE1 cells, whereas removal of I-helix further promoted condensate formation (Figs. S3F-I).

**Fig. 3.**
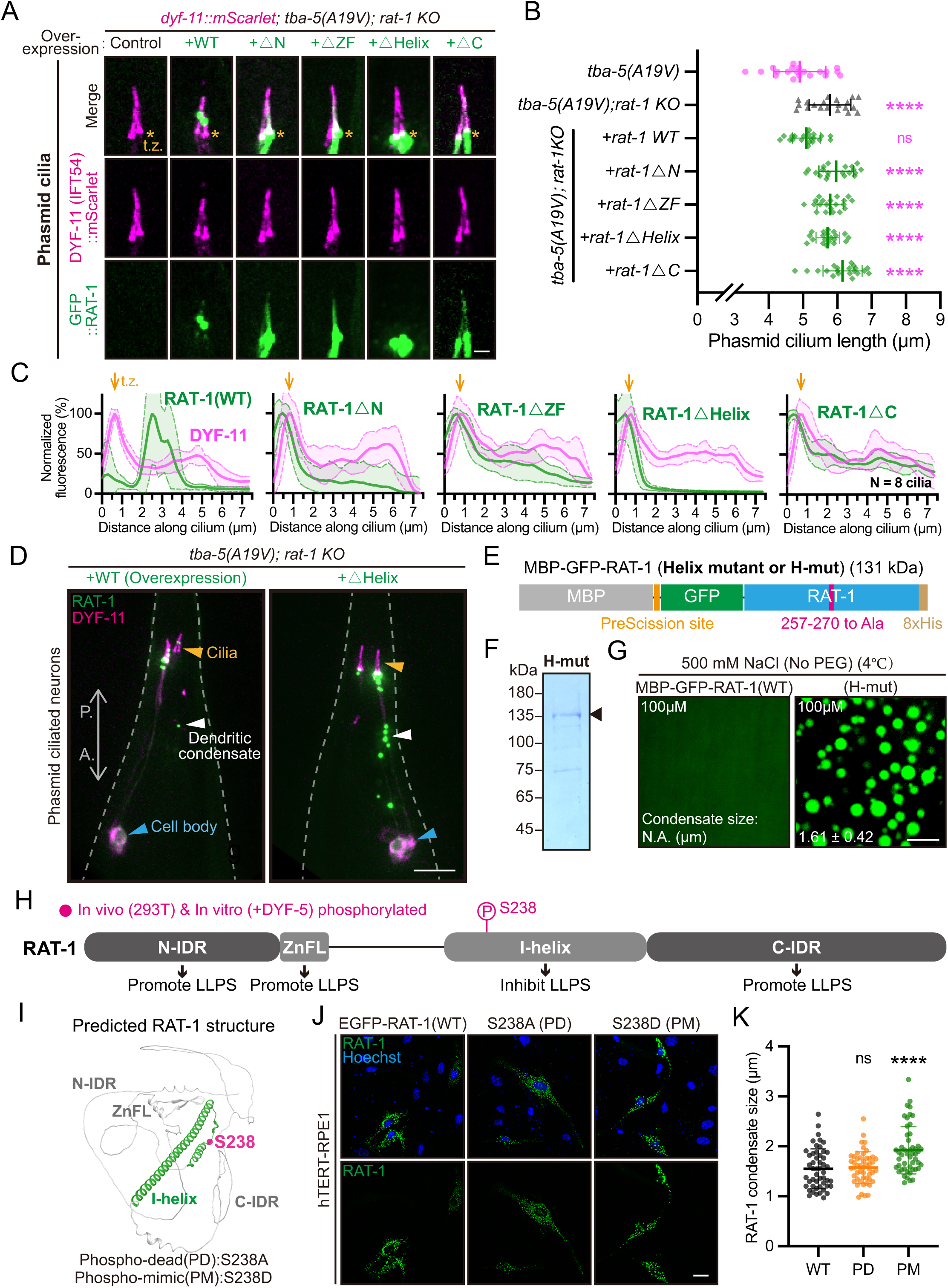
Phosphorylation regulates the internal α-helix (I-helix) for effective RAT-1 condensation. (A) Representative images of phasmid cilia in *tba-5 (A19V); rat-1 KO* animals overexpressing wild-type GFP::RAT-1 or truncation mutants. Cilia are marked with DYF-11(IFT54)::mScarlet (knock-in). t.z., transition zone (indicated by orange asterisks). Scale bar, 2 μm. (B) Quantification of phasmid cilia length in each group in (A). N = 20 phasmids. Shown as mean ± SD. One-way ANOVA with Dunnett’s tests were performed. (C) Distribution of RAT-1 and DYF-11 fluorescence along the cilium for each group in (A). 8 phasmid cilia were used for quantification and the average fluorescence was shown (mean ± SD). The maximal fluorescence in each group was normalized to 100%. (D) Representative images of phasmid ciliated neurons in *tba-5 (A19V); rat-1 KO* animals overexpressing WT GFP::RAT-1 or the △Helix mutant. Cell bodies of the ciliated neurons are indicated in blue, dendritic RAT-1 condensates in white, and cilia in orange. A., anterior; P., posterior. Scale bar, 10 μm. (E) Schematic of recombinant MBP-GFP-RAT-1 (Helix mutant or H-mut) for in vitro purification. See also Fig. S3J for more details. (F) SDS-PAGE analysis showing purification of MBP-GFP-RAT-1 (H-mut). (G) Representative images showing biophysical behaviors of 100 μM MBP-GFP-RAT-1 (WT or H-mut) in the presence of 500 mM NaCl at 4 ℃ (without PEG-8,000). Quantification of condensate size was shown as mean ± SD (N = 20 condensates from 3 independent experiments). Scale bar, 5 μm. (H) Mass spectrometry-identified in vivo (in HEK293T cells) and in vitro (treated with 0.5 μM *Ce*DYF-5) phosphorylation sites along the RAT-1 sequence. LLPS, liquid-liquid phase separation. (I) Predicted RAT-1 structure highlighting the internal helix (I-helix, green) and S238 (magenta). (J) Representative images showing overexpressed EGFP-RAT-1 (WT, PD or PM mutant) forms liquid-like condensates in hTERT-RPE1 cells. Scale bar, 10 μm. (K) Quantification of RAT-1 condensate sizes in each group in (J). N = 50 condensates from 3 independent experiments. Shown as mean ± SD. One-way ANOVA with Dunnett’s tests were performed. ns, not significant; **** *P* < 0.0001.

To further explore the inhibitory role of the I-helix, we generated a helix mutant with alanine substitutions at 14 central residues (L257-K270), predicted to disrupt the helix structure (Figs. 3E-F; Fig. S3J). This mutant underwent spontaneous phase separation even in high-salt conditions (500 mM NaCl) (Fig. 3G), indicating that the I-helix serves as an autoinhibitory element that suppresses premature condensate formation prior to ciliary entry. We also mutated four electropositive residues (K267-K270) within the I-helix, which significantly enhanced condensate formation in human RPE1 cells (Figs. S4A-B). Charge-neutralizing single mutations had milder effects, suggesting a synergistic role of these residues in inhibiting condensate formation. Interestingly, the isolated I-helix did not undergo phase separation on its own (Fig. S4C) but effectively suppressed condensate formation in the RAT-1 (△Helix) mutant when expressed separately (Figs. S4D-E). These results indicate that the I-helix inhibits condensation through interactions with other RAT-1 modules, rather than through structural confinement alone.

### Phosphorylation spatially regulates condensate formation by RAT-1

The regulation of RAT-1 condensate formation specifically at the distal ciliary regions has remained unclear. To investigate whether phosphorylation plays a role, we focused on DYF-5, a ciliary kinase in *C. elegans* that localizes to the distal ciliary regions, similar to RAT-1 (Fig. S5A) (Burghoorn et al., 2007; Li et al., 2021). In vitro kinase assays demonstrated that DYF-5 directly phosphorylates RAT-1 (Figs. S5B-C), enhancing its condensate formation (Figs. S5D-E). Mass spectrometry analysis in human HEK293T cells and DYF-5-mediated in vitro kinase assays identified Serine 238 (S238) within the I-helix as the potential phosphorylation site (Figs. 3H-I; Fig. S5F). Phospho-mimic (PM) mutant (S238D) enhanced RAT-1 condensate formation in human RPE1 cells, while phospho-dead (PD) mutant (S238A) displayed behavior similar to wild-type (Figs. 3J-K; Fig. S5G). These results suggest that phosphorylation at S238 relieves the autoinhibitory effect of the I-helix on phase separation.

To dissect the functional contribution of S238 phosphorylation to RAT-1 physiological activity, we generated GFP-tagged RAT-1 PD mutant (S238A) and PM mutant (S238D), and expressed them in *tba-5 (A19V); rat-1* knockout animals (Fig. S5H). While wild-type RAT-1 effectively reversed (reduced) the ciliary length in double-mutant animals, both PD and PM mutants conferred only mild effects on ciliary length (Fig. S5I), indicating partial loss of function. Specifically, the PD mutation attenuated phase separation capacity of RAT-1 and decreased the number of RAT-1 condensates within primary cilia (Fig. S5I). In contrast, the PM mutation prevented ciliary entry of RAT-1 and triggered premature condensate formation in dendrites, mirroring the phenotype observed for the RAT-1 (△Helix) mutant (Fig. S5H). These results imply that spatially confined phosphorylation (potentially on the S238 site) enables the physiological functionality of RAT-1.

Taken together, our findings support a model in which the I-helix acts as a gatekeeper, preventing premature condensation during dendritic transport (Fig. 4A). Site-specific regulation at or near S238 is required for proper RAT-1 function, as both phospho-dead and phospho-mimic mutants impair its activity in vivo. We therefore propose that spatially restricted phosphorylation or other post-translational modification(s) may regulate RAT-1 condensation at the ciliary tip, although the identity of the responsible kinase(s) (e.g. DYF-5) and the exact nature of the modification remain to be determined.

**Fig. 4.**
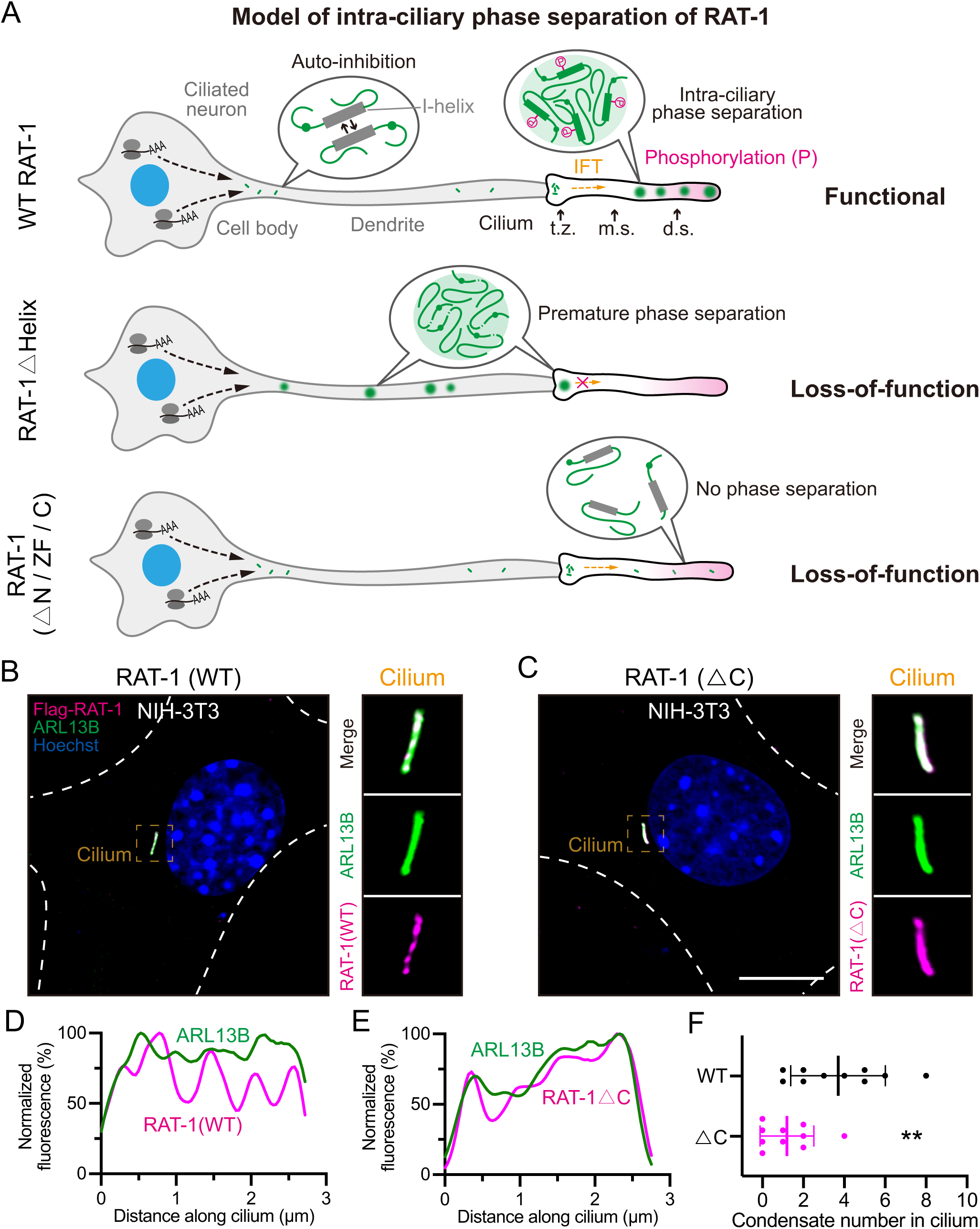
RAT-1 forms condensates within the cilia of mammalian cells. (A) Schematic models showing a graded control of the intra-ciliary phase separation behavior of RAT-1. For wild-type (WT) RAT-1 (top), an internal helix (I-helix)-mediated auto-inhibition mechanism prohibits the premature condensate formation in dendrites, which guarantees the ciliary entrance of RAT-1 through intraflagellar transport (IFT). RAT-1 may be phosphorylated in the distal region of cilia, further promoting the intra-ciliary condensate formation (phase separation). For the △Helix mutant (middle), RAT-1 undergoes premature condensate formation in dendrites, which abolishes its ciliary entrance. For the △N-IDR/ZnFL/C-IDR mutant (bottom), RAT-1 fails to undergo intra-ciliary phase separation and is loss-of-function. t.z., transition zone; m.s., middle segment; d.s., distal segment. (B-C) Representative immunofluorescence images showing the localization of transfected wild-type RAT-1 (B) or the △C-IDR mutant (C) in mammalian cilia (NIH-3T3 cell line). ARL13B marks the cilia. Scale bar, 10 μm. (D) Distribution of RAT-1 and ARL13B fluorescence along the cilium in (B). The maximal flurescence was normalized to 100%. (E) Distribution of RAT-1 and ARL13B fluorescence along the cilium in (C). (F) Quantification of RAT-1 (WT or △C-IDR) condensate numbers in the NIH-3T3 cilia. N = 10 cilia. Shown as mean ± SD. Nonparametric (Mann-Whitney) tests were performed. ** *P* < 0.01.

### RAT-1 exhibits condensation behavior within mammalian cilia

To investigate whether the condensate-forming properties of RAT-1 extend beyond *C. elegans*, we expressed Flag-tagged worm RAT-1 in the NIH-3T3 mammalian ciliated cell line. At low expression levels, RAT-1 localized specifically to the cilia and formed discrete condensates, recapitulating the punctate distribution seen in worm sensory cilia (Fig. 4B). Condensate formation was dependent on the C-terminal intrinsically disordered region (C-IDR), as deletion of this region (△C) abolished the ciliary puncta without affecting overall protein expression or ciliary entry (Figs. 4C-F; Figs. S5J-L). These findings suggest that RAT-1’s ability to form spatially confined condensates may be a feature that could also be relevant for regulating ciliary MT dynamics in other species.

### RAT-1 promotes tubulin turnover at the ciliary tip

Given the tubulin-specific interaction between *rat-1* and *tba-5*, we investigated whether RAT-1 regulates tubulin dynamics within cilia. Using the split-GFP system to label endogenous TBA-5 (Fig. S1E), we performed FRAP analysis at the ciliary tip (Figs. 5A-B). Previous work has shown that both wild-type TBA-5 and the TBA-5 (A19V) mutant undergo active turnover at ciliary tips, with the mutant displaying altered incorporation dynamics associated with disruption of ciliary distal segments (Xu et al., 2024). Consistent with these findings, TBA-5 (WT) in a wild-type *rat-1* background displayed rapid fluorescence recovery at the ciliary tip, indicating dynamic tubulin turnover (Figs. 5A-D). In contrast, *rat-1* knockout animals showed markedly reduced recovery for both wild-type and mutant TBA-5 (A19V) (Figs. 5A-D), indicating a severe impairment in tubulin turnover.

**Fig. 5.**
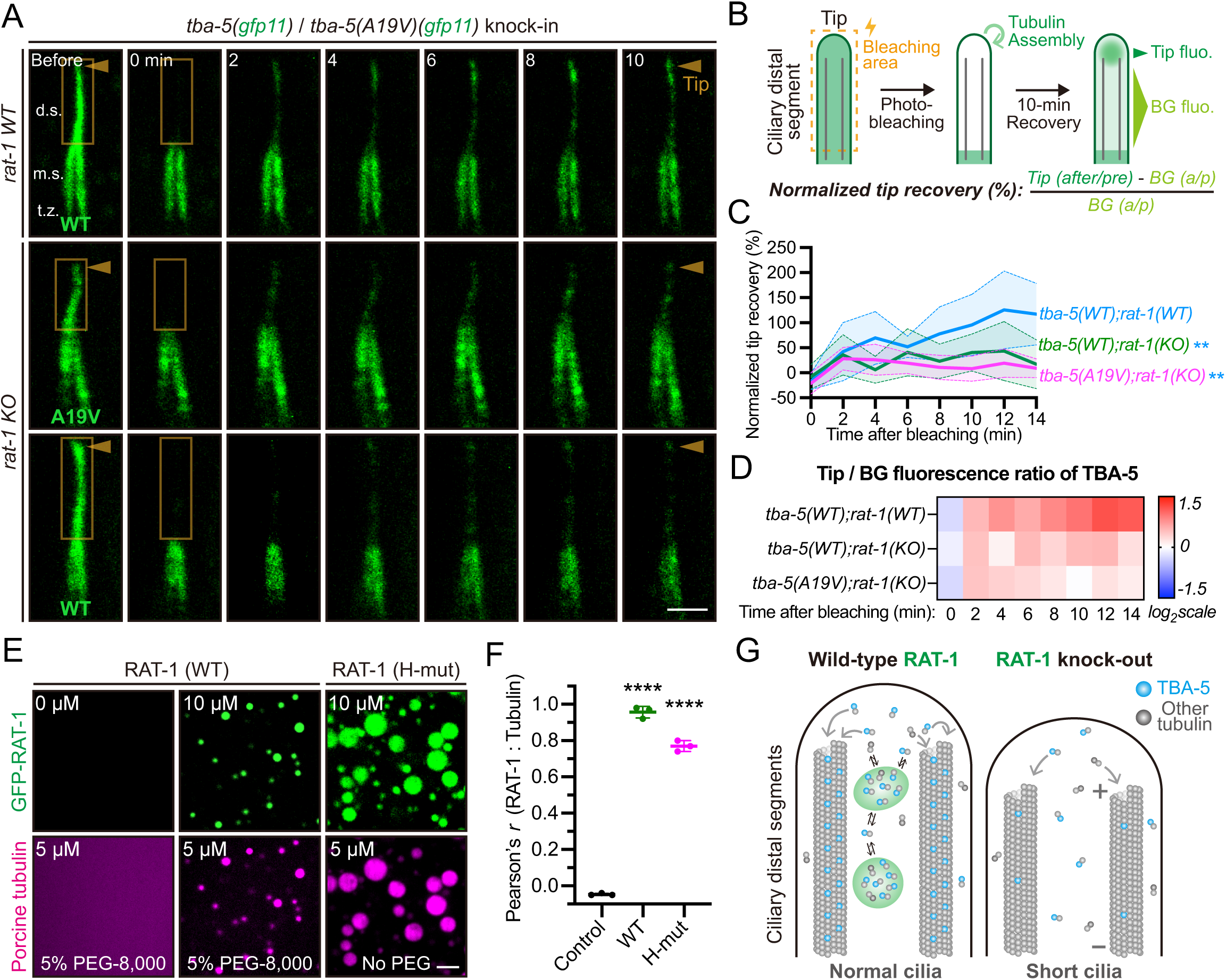
RAT-1 condensates promote tubulin turnover at ciliary tips. (A) Representative FRAP (fluorescence recovery after photobleaching) images of split-GFP-labeled TBA-5 (WT) or TBA-5 (A19V), in *rat-1* WT or KO background. Ciliary tips are indicated by orange arrowheads. Bleaching zones are indicated by orange boxes. t.z., transition zone; m.s., middle segment; d.s., distal segment. Scale bar, 2 μm. (B) Schematic model of the fluorescence quantification of GFP11-i-labeled tubulins after photobleaching. Ciliary distal segments are bleached specifically, followed by recovery in dark. The fluorescence intensities of ciliary tips (Tip fluo.) or distal segments except for the tips (BG fluo.), are measured before or after the bleaching. (C) Normalized tip recovery in each group in (A). Shown as mean ± 95 % CI. N = 10 cilia. Unpaired t tests were performed at 14 min after bleaching. ** *P* < 0.01. (D) Heatmap representation of Tip/BG fluorescence ratio at each time after bleaching. N = 10 cilia. Data are presented in log2 scale. (E) Representative images showing in vitro localization of rhodamine-labeled porcine brain tubulins in condensates formed by GFP-RAT-1 (WT or H-mut). MBP tag was removed. Scale bar, 5 μm. (F) Pearson’s *r* between RAT-1 and tubulin fluorescence in each group in (E). 3 independent replicates were performed per group. ‘Control’ represents the tubulin only group. One-way ANOVA with Dunnett’s tests were performed. **** *P* < 0.0001. (G) A proposed model showing that RAT-1 promotes tubulin turnover at ciliary tips. RAT-1 forms intra-ciliary condensates and may serve as the tubulin reservoir, which promotes ciliary tubulin dynamics (such as incorporation into ciliary microtubules).

These observations imply that RAT-1 may interact with ciliary tubulins in vivo. To investigate this, we transgenically expressed GFP::RAT-1 and TBA-5::mScarlet in wild-type animals. Within the amphid cilia, GFP::RAT-1 effectively enriched TBA-5::mScarlet into the condensates (Fig. S6A), exhibiting partial co-localization in cilia (Fig. S6B, Pearson’s *r* = 0.63 ± 0.11). After photobleaching of TBA-5 fluorescence followed by recovery for 90 seconds, we detected enriched recovery of TBA-5 fluorescence within the RAT-1 condensates (Figs. S6A-B), indicating that RAT-1 condensates dynamically enrich and interact with TBA-5. Consistent with these in vivo findings, GFP-RAT-1 condensates also co-localized with rhodamine-labeled porcine brain tubulins in vitro (Figs. 5E-F), further supporting a direct interaction between RAT-1 and tubulin. Additionally, in *tba-5 (A19V)* mutants, which exhibit temperature-sensitive ciliary defects (Figs. S6C-E), GFP::RAT-1 formed discrete condensates near the ciliary middle segment tip, regardless of ciliary length, further supporting its role in promoting tubulin turnover. Together, these findings suggest that RAT-1 condensates locally enrich tubulin and are associated with its dynamic exchange at the ciliary tip (Fig. 5G).

Given that the TBA-5 (A19V) mutant impairs ciliary MT function in a toxic, ‘gain-of-function’ manner, and loss of TBA-5 results in recovered ciliary distal segment and dye-filling phenotype (Hao et al., 2011), we propose that loss of RAT-1 suppresses TBA-5 (A19V) toxicity by reducing the incorporation and turnover of mutant tubulin within ciliary MTs. Consistent with this model, loss of RAT-1 decreases tubulin dynamics at ciliary tips (Figs. 5A-B) and partially phenocopies reduced TBA-5 function, including ivermectin resistance (Fig. S1H). Although this mechanism remains to be tested directly, it provides a plausible explanation for the suppressor effect of *rat-1* mutations.

### RAT-1 interacts with EB family proteins to support MT stability

Although RAT-1 forms condensates and enriches tubulins (Fig. 2C), its interaction with the MT plus ends, where ciliary MTs polymerize, remains unclear. Interestingly, we observed that the localization of RAT-1 condensates in wild-type animals is temperature-dependent. At 15 °C, RAT-1 condensates were dispersed within ciliary distal regions (Fig. S7A), while at 25 °C, they accumulated at the tips of both the middle and distal ciliary segments (Fig. S7B). Time-lapse imaging further revealed that RAT-1 translocates toward the ciliary tips as the temperature increases (Figs. S7C-D), coinciding with the enriched localization of EB family proteins at these ciliary tips (Hao et al., 2011). Indeed, GFP::RAT-1 partially colocalized with EBP-2::mScarlet (the homolog of human EB) at the ciliary tips in *C. elegans* sensory neurons (Fig. 6A).

**Fig. 6.**
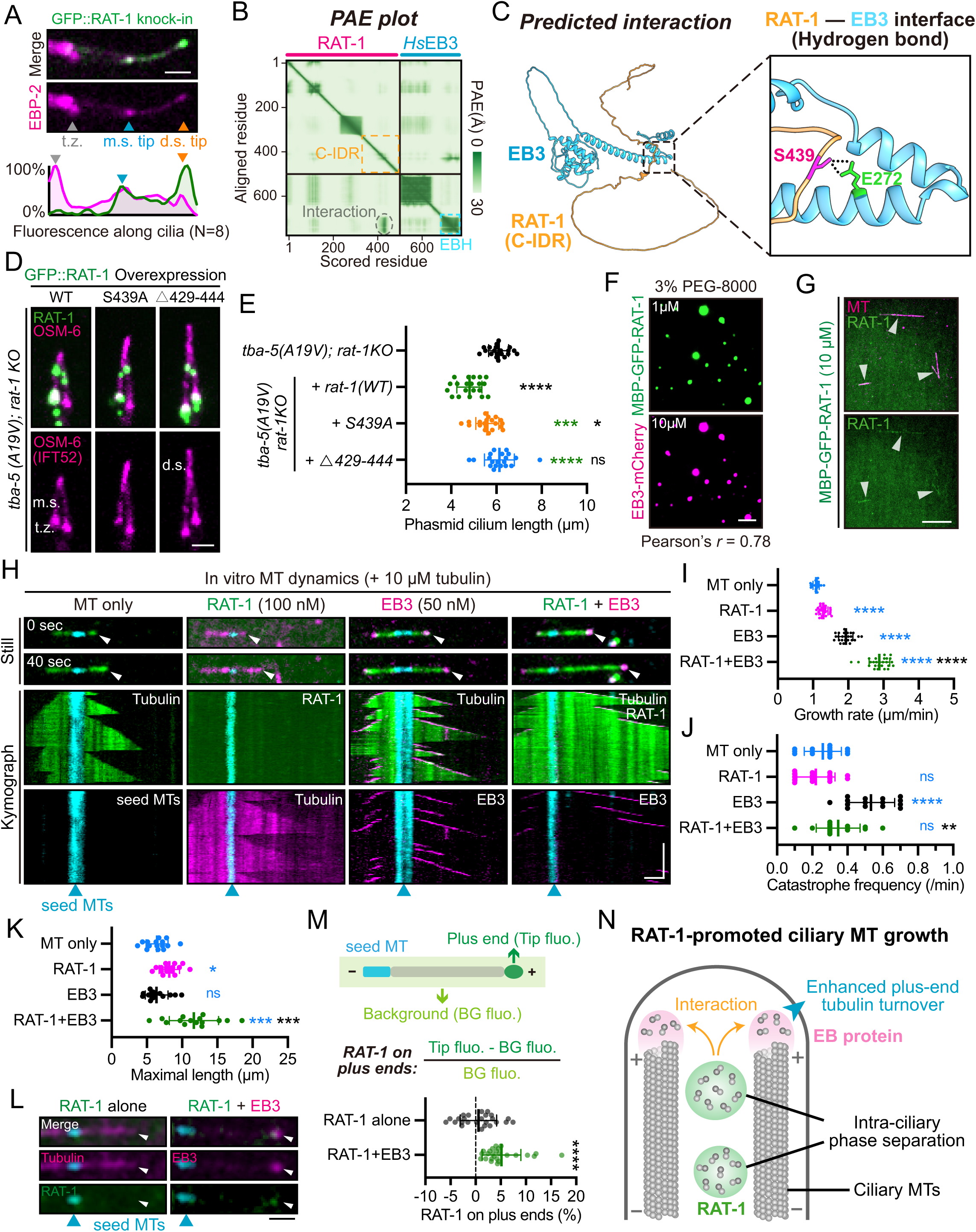
RAT-1 cooperates with end-binding (EB) proteins to promote microtubule growth. (A) (Top) Representative images of *C. elegans* phasmid cilia with GFP::RAT-1 (knock-in). EBP-2::mScarlet (human EB homolog) was overexpressed as a marker of ciliary microtubule plus ends, corresponding to the tips of middle segments (m.s.) and distal segments (d.s.). (Bottom) Average fluorescence distribution of RAT-1 (green) and EBP-2 (magenta) along the phasmid cilia (N = 8). The minimal and maximal fluorescence intensities were normalized to 0 % and 100 %, respectively. t.z., transition zone. Scale bar, 2 μm. (B) AlphaFold3-predicted PAE (predicted alignment error) plot of the RAT-1 and *H. sapeins* (*Hs*) EB3 interaction. C-IDR was indicated in orange. EB homology domain (EBH) was indicated in blue. Expected position error (Å) ranges from 0 (green) to 30 (white). (C) Predicted interaction pattern of the RAT-1 C-IDR (orange) with EB3 (blue). Key interacting residues are marked in magenta (RAT-1 S439) or green (EB3 E272). Hydrogen bonds are indicated by dotted lines. (D) Representative images of phasmid cilia in *tba-5 (A19V); rat-1 KO* animals overexpressing wild-type GFP::RAT-1, or S439A, or △429-444 mutant. Cilia are marked with OSM-6(IFT52)::mScarlet (overexpression). Scale bar, 2 μm. (E) Quantification of phasmid cilia length in each group in (D). N = 20 phasmids. Shown as mean ± SD. One-way ANOVA with Tukey’s tests were performed. (F) Representative images showing in vitro co-phase separation of MBP-GFP-RAT-1 and EB3-mCherry (Pearson’s *r* = 0.78 from 3 independent replicates). Scale bar, 5 μm. (G) Representative in vitro images showing co-localization of MBP-GFP-RAT-1 and rhodamine-conjugated polymerized MTs. Scale bar, 10 μm. (H) Representative TIRF (total internal reflection fluorescent microscope) images of in vitro microtubule (MT) dynamics assay with 10 μM tubulins (+ 1% PEG-8,000). Kymographs display the time-lapse dynamics of growing MTs. HiLyte647-labeled seed MTs were employed for all groups. For MT only / EB3 / RAT-1+EB3 group, HiLyte488-labeled tubulins (green) were employed. For RAT-1 group, rhodamine-labeled tubulins were employed. Growing MT plus ends are indicated by white arrowheads. Scale bar, 2 μm (horizontal) and 2 min (vertical). See also Movie S3. (I-K) Quantification of MT growth rate (N = 25, Welch’s ANOVA with Dunnett’s T3 tests) (I), catastrophe frequency (N = 15, Kruskal-Wallis with Dunn’s tests) (J) and maximal growth length (N = 15, Welch’s ANOVA with Dunnett’s T3 tests) (K) for each group in (H). Data were collected from three independent MT dynamics assays. N indicates the number of individual MTs analyzed across all experiments. Measurements were obtained during a 10-min growth period. ns, not significant; * *P* < 0.05; ** *P* < 0.01; *** *P* < 0.001; **** *P* < 0.0001. See also Movie S3. (L) Representative TIRF images showing localization of MBP-GFP-RAT-1 (100 nM) at growing MT plus ends (indicated by white arrowheads), in the absence (RAT-1 alone) or presence of 50 nM EB3-mCherry (RAT-1+EB3). MT dynamics assays were performed with 10 μM rhodamine-conjugated tubulins (RAT-1 alone) or unlabeled tubulins (RAT-1+EB3) (+ 1% PEG-8,000). Seed MTs were indicated by blue arrowheads. Scale bar, 1 μm. (M) (Top) Schematic illustrating quantification of relative RAT-1 amounts on growing MT plus ends. RAT-1 fluorescence (green) on MT plus ends was measured as Tip fluo., and background fluorescence was measured as BG fluo.. (Bottom) Quantification of RAT-1 intensities on dynamic MT plus ends in the absence or presence of EB3. N = 25 independent MTs from 3 independent assays. Shown as mean ± SD. Unpaired t test was performed. (N) A proposed model illustrating RAT-1-promoted ciliary MT growth. RAT-1 undergoes intra-ciliary phase separation. Through interaction with plus end-binding (EB) proteins, RAT-1 promotes local tubulin turnover, as well as MT growth.

This prompted us to investigate the potential interaction between RAT-1 and EB family proteins. AlphaFold3 predictions identified a conserved interface between RAT-1’s C-IDR and EB family proteins, including human EB1, EB2, and EB3, with a key hydrogen bond between RAT-1 S439 and EB3 E272 (Figs. 6B-C; Figs. S7E-G). Mutants disrupting this interaction, such as S439A and △429-444, failed to rescue ciliary defects in *tba-5 (A19V); rat-1* knockout animals (Figs. 6D-E; Fig. S7H), though they retained the ability to form condensates (Fig. S7I). In vitro pull-down assays demonstrated that RAT-1 directly interacts with EB3, and deletion of residues 429-444 markedly weakened this interaction (Figs. S7J-K). Additionally, RAT-1 and EB3 formed co-phase-separated droplets (Fig. 6F), with enhanced interaction at elevated temperatures (Figs. S7L-M), consistent with the in vivo observation of temperature-dependent RAT-1 translocation (Figs. S7A-D).

We next assessed RAT-1’s interaction with polymerized MTs. In vitro co-incubation assays revealed that RAT-1 (10 μM) showed a modest affinity for rhodamine-labeled MTs (Fig. 6G). MT dynamics assays demonstrated that RAT-1 alone (100 nM) slightly but significantly increased MT growth rates with minimal effects on catastrophe frequency, resulting in a mild enhancement of MT elongation (Figs. 6H-K; Movie S3). EB3 alone (50 nM) increased both MT growth rates and catastrophe frequency, consistent with previous studies (Figs. 6H-K; Movie S3) (Bieling et al., 2007). When combined, RAT-1 (100 nM) and EB3 (50 nM) acted synergistically to promote persistent MT growth: catastrophe frequency was markedly reduced, growth rates were further enhanced, and maximal microtubule extension exceeded that observed with either protein alone (Figs. 6H-K; Movie S3).

These findings indicate that RAT-1 not only enriches tubulin within condensates but also cooperates with EB family proteins to stabilize MTs and promote efficient elongation. To examine whether EB proteins influence the localization of RAT-1 at growing MT ends, we performed time-lapse imaging of dynamic MTs. EB3 exhibited comet-like localization at growing MT plus ends, and RAT-1 showed enhanced enrichment at EB3-positive tip regions (Fig. 6L). Notably, this plus-end association of RAT-1 was rarely observed in the absence of EB3 (Figs. 6L-M). Collectively, these results show that RAT-1 acts as the local hub to concentrate tubulin and cooperate with EB family proteins, providing a mechanism for sustained MT growth at the ciliary tip (Fig. 6N).

## Discussion

In this study, we establish RAT-1 as a regulator of ciliary MT dynamics and tubulin homeostasis. The ability of RAT-1 condensates to enrich tubulin, together with the altered tubulin dynamics observed in *rat-1* knockout mutants, suggests that RAT-1 may function as a local tubulin reservoir within cilia. Moreover, RAT-1 cooperates with EB family proteins to stabilize MTs and prevent catastrophe, ensuring efficient MT elongation. Our results offer insights into the spatially restricted regulation of ciliary MT growth, which is essential for maintaining ciliary structure and function.

Recent studies have shown that EB1 and its family members undergo phase separation to facilitate their function at MT plus ends (Maan et al., 2023; Meier et al., 2023; Song et al., 2023), enhancing MT polymerization and stability. Our study extends these findings by revealing an additional mechanism in which RAT-1 forms liquid-like condensates that not only localize tubulin to the ciliary tip but also directly cooperate with EB3 to stabilize MTs and promote their elongation. While EB proteins primarily function at MT plus ends, RAT-1 may contribute to coordination between tubulin supply and MT dynamics through spatially regulated phase separation, providing a more precise mechanism for local tubulin enrichment and MT growth.

This work advances our understanding of phase separation by linking it to both tubulin turnover and EB function at specific subcellular locations, offering a unique framework for orchestrating MT regulation in confined environments like cilia. While previous studies have highlighted phase separation in processes like chromatin remodeling during cell division and RNA processing in stress granules (Molliex et al., 2015; Sanulli et al., 2019), its role in regulating MTs—particularly in the highly specialized structure of cilia—has remained largely unexplored. By locally concentrating tubulin at the ciliary tip, RAT-1 condensates facilitate efficient tubulin turnover and incorporation into growing MTs, thereby enhancing our understanding of how cells utilize phase separation to precisely regulate MT polymerization and stabilization in dynamic, subcellular environments.

The mechanisms uncovered in this study have broad implications for understanding MT dynamics in tightly confined cellular compartments like cilia. Phase separation of tubulin-binding proteins, such as RAT-1, provides a model for how cells locally concentrate and regulate essential structural components in specific subcellular locations. This concept aligns with previous work on BuGZ (Jiang et al., 2015), which demonstrated tubulin co-phase separation as a mechanism for MT organization in mitotic spindles. Although we did not identify direct human homologs of RAT-1 due to its largely unstructured nature, proteins like BuGZ that also undergo phase separation with tubulin share similar intrinsically disordered regions (IDRs) and α-helical elements, suggesting that RAT-1 may operate via a comparable mechanism. This highlights a potential common mechanism for MT regulation via phase separation across different protein families, even in the absence of sequence homology. Furthermore, the ability of RAT-1 to form condensates in mammalian cells and localize to mammalian cilia supports the idea that this behavior is evolutionarily conserved.

Mechanistically, the biophysical behavior of RAT-1 is tightly controlled by an internal α-helical region (I-helix), which prevents premature condensation during dendritic transport, and by phosphorylation from the ciliary kinase DYF-5/MAK (Burghoorn et al., 2007; Jiang et al., 2022), which locally relieves this inhibition to trigger distal region-specific condensate formation. This combination of intrinsic autoinhibition and spatially restricted kinase activity ensures that tubulin is concentrated precisely where it is needed, coupling material supply to growth while preventing ectopic condensation elsewhere. Such spatial confinement provides a conceptual framework for understanding how phase separation can be harnessed in small, crowded organelles to regulate cytoskeletal assembly.

Functionally, this work investigates long-standing questions regarding how ciliary tubulins are maintained within cilia after unloading from the IFT complex (Jiang et al., 2022), and whether a membrane-less reservoir exists within cilia for local material enrichment and supply (Johnson and Malicki, 2019). More broadly, the kinase-regulated, spatially confined phase separation described here may represent a general strategy by which cells coordinate polymer assembly within small organelles or specialized compartments.

This study has several limitations. While the condensate formation behavior of RAT-1 is facilitated by phosphorylation, additional molecular mechanisms and post-translational modifications may also have regulatory roles within cilia. The mild ciliary defects (short cilia and partial resistance to ivermectin) observed in *rat-1* knockout worms suggest redundancy or compensation by other tubulin regulators, raising the possibility that additional phase-separated ciliary proteins remain to be discovered. In addition to locally concentrating tubulin, RAT-1 could potentially influence tubulin stability, trafficking, folding, or the activity of other MT-associated proteins. Furthermore, how RAT-1 cooperates with EB proteins and whether additional MT regulators participate in this process remain important questions for future studies.

In summary, our work reveals that RAT-1 forms kinase-regulated, spatially confined condensates at the ciliary distal region to locally concentrate tubulin and coordinate MT growth. An internal α-helix (I-helix) prevents its premature condensation during transport, while phosphorylation licenses assembly precisely where it is needed. RAT-1 cooperates with end-binding proteins to suppress catastrophe and enhance MT elongation, providing a mechanism that couples material supply to cytoskeletal dynamics. These findings uncover a previously unrecognized strategy by which phase separation organizes polymerization within confined cellular compartments, offering new insights into ciliary biology and the broader principles of cytoskeletal regulation.

## Materials and Methods

### Worm strains and genetic cross

*C. elegan*s were cultured at 20 ℃ on nematode growth medium (NGM) plates with the *Escherichia coli* OP50 unless described otherwise. For genetic cross, 20 μL suspended OP50 was dropped at the center of NGM plates to make a cross plate, and 10 males carrying *him-5 (e1490)* allele and 5 hermaphrodites were transferred to this plate for at least 24 hours. F1 and F2 hybrid progenies were screened by Sanger sequencing using EasyTaq® 2× Super Mix (TransGen Biotech, #AS111-14). Dataset S1 summarizes the strains used in this study.

### Molecular biology

For over-expression (OE) plasmids in *C. elegans*, cDNA construct of *F41E7.9 (rat-1)*, or genome DNA construct of *Prat-1::rat-1* containing 550 bp upstream promoter and 300 bp downstream 3’UTR (untranslated region), was cloned into pDONR vector. For knock-out (KO) and knock-in (KI) plasmids, 20 bp CRISPR-Cas9 target (‘ACTGGATGCATTGAAAGCGT’ in the first exon of rat-1) was inserted into the pDD162 vector (Addgene #47549) by linearizing this vector with 20 bp overlapped primers. The resulting PCR products were treated with DpnI digestion for 4 hours and then transformed into *E. coli* Trans5α. The linearized PCR products were cyclized by spontaneous recombination in bacteria. The homology recombination (HR) templates were constructed by cloning the 1.0 kb upstream and downstream homology arms into pPD95.77 vector (Addgene #37465) using In-Fusion® HD Cloning Kit (Takara Bio, #639650). Ultimately, CRISPR-Cas9-targeted sites in the templates were modified with synonymous mutations. For mammalian cell transfection, cDNA construct of nematode *rat-1* and its truncation mutants were cloned into pCMV3.0 vector containing CMV enhancer, CMV promoter, EGFP and bGH terminator. For purification of recombinant RAT-1 from bacteria, cDNA of maltose binding protein-PreScission site-GFP-8xHis (MBP-3C-GFP-His), or MBP-3C-GFP-RAT-1-His was cloned into pET21 vector, downstream of T7 promoter (Addgene #89881). All plasmids were purified with AxyPrep Plasmid Purification Miniprep Kit (Axygen, #AP-MN-P-250) and PureLink Quick PCR purification Kit (Invitrogen, #K310001).

### Extra-chromosomal transgenesis (over-expression) and genome editing

Transgenic lines of *C. elegans* were obtained by co-injecting the over-expression (OE) plasmids with *rol-6 (su1006)* marker (20 ng/μL) into the gonads of young adult worms. The concentrations for micro-injection are: *Prat-1::rat-1* (5 ng/μL), *Pdyf-1::gfp::rat-1* (5 ng/μL), *Pdyf-1::ebp-2::mscarlet* (10 ng/μL), *Pdyf-1::osm-6::mscarlet* (10 ng/μL), *Pdyf-1::tba-5::mscarlet* (20 ng/μL). The concentrations of mutated constructs were consistent with their wild-type forms. At least two independent transgenic lines with a constant transmission rate (> 50%) were examined and analysed. CRISPR-Cas9-mediated genome editing was used to create knock-out (KO) or knock-in (KI) strains (Cong et al., 2013). Target sequences were selected by the CRISPR design tool (https://crispor.gi.ucsc.edu). For knock-out, CRISPR-Cas9 construct with target was co-injected into the gonads of young adult worms at 20 ng/μL with *rol-6 (su1006)* (20 ng/μL). For knock-in, the CRISPR-Cas9 constructs with targets and HR templates were co-injected at 50 ng/μL with *rol-6 (su1006)* (50 ng/μL). F1 transgenic progenies (roller) were singled and screened by PCR. All knock-out and knock-in alleles were verified by the Sanger sequencing of the entire genes to guarantee that no other mutations were introduced. SunyBiotech Ltd. generated some knock-in strains and alleles in this study. The name of these strains started with ‘PHX’; The name of these alleles started with ‘syb’.

### Dye-filling assay

The fluorescence dye DiI (1,1′-dioctadecyl-3,3,3′,3′-tetramethylindocarbocyanine perchlorate, Sigma) and dye-filling assay was broadly used to assess the ciliary function (Inglis et al., 2007; Perkins et al., 1986). Dye-filling positive animals possess relatively intact ciliary structures, while dye-filling defective animals develop abnormal cilia. Worms with *tba-5 (A19V)* mutant background were cultured at 15 ℃ for at least 12 hours before this assay. Young adult worms were harvested into 500 µL M9 buffer with DiI (4 µg/ml), followed by incubation at 15 ℃ in the dark for 1 hour. Worms were then transferred to seeded NGM plates and examined for dye uptake 2 hours later using a fluorescence compound microscope. We observed at least 100 animals of each strain from three independent assays.

### Ivermectin (IVM) resistance assay

Ivermectin is widely used to treat numerous parasitic infections of humans and livestock (Campbell, 1993). *C. elegans* cilia mutants display IVM resistance, primarily due to defective IVM uptake by amphids (Dent et al., 2000; Page, 2018). For IVM resistance assay, synchronized L1 (larvae 1) worms were transferred to NGM plates containing 10 ng/mL IVM. After culturing at 20 ℃ for 3 days, percentage of larvae arrested worms (fail to reach young adult stage) was counted.

### FRAP (fluorescence recovery after photo-bleaching)

FRAP experiments were carried out using Zeiss LSM900 with Airyscan2 confocal microscopy (Carl Zeiss). In *C. elegans*, *gfp::rat-1* knock-in strain was employed for FRAP experiments of RAT-1 condensates in cilia at 20 ℃. A 488 nm laser at 100 % power was used for photobleaching, and images were acquired every second (half-bleach) or 3 sec (full-bleach). *tba-5 (gfp11-i)* or *tba-5 (A19V) (gfp11-i)* knock-in worms with *che-3::t2a::gfp1-10* were employed for FRAP experiments of TBA-5 in ciliary distal segments at 15 ℃. Images were acquired every 2 min after bleaching. We measured fluorescence intensities at the ciliary tip (Tip) or distal segments except for tip (BG) before or after the bleach. To calculate normalized tip recovery, tip recovery ratio (Tip after bleaching divided by Tip before bleaching) is subtracted by BG recovery ratio (BG after bleaching divided by BG before bleaching), and then divided by BG recovery ratio. *gfp::rat-1; tba-5::mscarlet* overexpression strain was employed for FRAP experiments of TBA-5 in cilia at 20 ℃. A 561 nm laser at 100 % power was used for photobleaching, and images were acquired every 15 sec. For in vitro experiments, 10 μM GFP-RAT-1 (with MBP tag removed through co-incubation with 1 x HRV 3C protease (Takara) for 30 min at room temperature) was incubated with 5 % PEG-8000 at 20 ℃ to initiate liquid-like phase separation (LLPS). After bleaching, images were acquired every 5 sec.

### Cell culture and transfection

HEK293T and NIH-3T3 cells were grown in DMEM medium (Gibco) containing 10 % fetal bovine serum (FBS) (Yeasen) and 1 % penicillin and streptomycin (Yeasen) at 37 ℃ with 5 % CO2. hTERT-RPE-1 cells were grown in DMEM/F12 medium (Procell) containing 10 % FBS and 1 % penicillin and streptomycin. Cells were seeded on 35 mm glass-bottom confocal dishes (CellVis, #D35C4-20-0-N) the day before transfection. Cells were transfected with 400 ng egfp-rat-1 per sample, as well as its truncation or other mutants. For transfection experiments involving the RAT-1 helix, cells were transfected with egfp-rat-1 (400 ng) + mcherry (600 ng, Mock) / egfp-rat-1 (400 ng) + mcherry-helix (600 ng) / egfp-rat-1△helix (400 ng) + mcherry (600 ng, Mock) / egfp-rat-1△helix (400 ng) + mcherry-helix (600 ng), respectively, using LipofectamineTM 3000 Transfection Reagent (Invitrogen) following manufacturer’s instruction. For immunofluorescence (IF) staining, cells were transfected with flag-rat-1 (400 ng) or flag-rat-1△C-IDR (400 ng).

### Live imaging of *C. elegans*

Worm hermaphrodites were anesthetized with 0.1 mmol/L levamisole in M9 buffer, mounted on 3 % agarose pads, and maintained at 20 ℃ unless described otherwise. Imaging was performed using an Axio Observer Z1 microscope (Carl Zeiss) equipped with 488 and 561 laser lines, a Yokogawa spinning disk head, an Andor iXon + EM-CCD camera, and a Zeiss 10×/0.25 or 100×/1.46 objective. Images were acquired at 0.8-μm z-stack (100×) or 5-μm z-stack (10×), or 200 / 700 ms intervals by µManager (https://www.micro-manager.org). Images were taken using identical settings (EM-gain: 250, exposure time: 200 ms, 10 % of max laser). For imaging of EBP-2::mScarlet, we used 100 % of max laser for improved resolution. Stacked images covering the whole cilia structures were used to measure the cilia length. Image analysis and measurement were performed with Fiji / ImageJ software (http://rsbweb.nih.gov/ij/). The visualized color of red channel was changed to colorblind-safe magenta in ImageJ. All images shown were adjusted linearly. Kymographs were generated in Fiji by manually drawing lines along cilia.

### Super-resolution imaging of mammalian cells

For mammalian cells, imaging was performed using Zeiss LSM900 with Airyscan2 confocal microscopy (Carl Zeiss) equipped with a 10×/0.45 or 63×/1.4 objective. Images were acquired at laser wavelength of 488 nm for EGFP/Dylight488, 561 nm for mCherry/Dylight549, 640 nm for Deep Red/Dylight649 and 405 nm for Hoechst. Images were taken using identical settings (Airyscan mode: Airyscan SR; Scan direction: bidirectional; Scan mode: frame). To induce ciliogenesis in NIH-3T3 or RPE-1 cells, the medium was replaced with serum-free opti-MEM medium (Gibco) after 8-h transfection, followed by 24-h culturing at 37 ℃ with 5 % CO2. The cells were incubated with DMEM medium containing 2 μg/mL Hoechst 33342 and 1x Tubulin TrackerTM Deep Red (Invitrogen, #T34076) at 37 °C for 30 min before live-cell imaging. For immunofluorescence (IF) staining, anti-DYKDDDDK first antibody (CST, #14793) (1:800 diluted) was used to stain flag-RAT-1; anti-ARL13B first antibody (Proteintech, #17711-1-AP and #66739-1-Ig) (1:400) was used to stain cilia. After incubation with first antibodies overnight at 4 ℃, cells were washed three times with PBS buffer. Secondary antibodies (Dylight 488/549/649, Abbkine) were added (1:300) for 1-hour incubation at RT, followed by washing three times with PBS. The cells were incubated with 1 μg/mL Hoechst 33342 for 10 min before imaging. All images shown were adjusted linearly. Fluorescence distribution was generated in Fiji by manually drawing lines along cilia.

### Purification of recombinant proteins

For expression and purification of recombinant MBP-3C(PreScission site)-GFP-His, MBP-3C-GFP-RAT-1-His, MBP-3C-GFP-RAT-1(H-mut)-His, StrepII-DYF-5 and EB3-mCherry-StrepII proteins, plasmids were transformed into BL21 (DE3) bacteria. 20-mL culture from a single colony was used to inoculate a 2-L fresh LB medium for further incubation at 37 °C with shaking. After 3.5-4 h, IPTG was added to the culture to a final concentration of 1 mM to induce protein expression at 18 °C for 16 h.

For RAT-1, bacteria pellets were collected by centrifugation and resuspended in 50 mL of ice-cold lysis buffer (20 mM HEPES, 1.2 M NaCl, 5 % glycerol, pH 8.2) containing 1 mM PMSF. After breaking bacteria by microfluidizer (AH-TITAN PLUS) at 800 bar for 15 min, the lysate was centrifuged at 12,000 rpm for 40 min at 4 °C. The supernatant was transferred to a fresh tube containing 4 mL of 50 % slurry of Ni-NTA beads (Smart-Lifesciences). After rotating at room temperature (RT) for 30 min, the mixture was transferred to a column to allow the settlement of the beads. The column was washed with 50 mL lysis buffer and 50 mL wash buffer (lysis buffer with 60 mM imidazole, pH 8.2), and eluted with 10 mL elution buffer (lysis buffer with 300 mM imidazole, pH 8.2) at RT. Eluates were concentrated to 50 μL with Amicon Ultra 30K device (Millipore) at 16 °C, as we found RAT-1 easily forms pellets (or condensates) below 8 °C. Imidazole was removed through buffer exchange during the ultrafiltration process.

For EB3, the lysis buffer was BRB80 (80 mM PIPES, 100 mM KOH, 1 mM EGTA, 1mM MgCl2, pH 6.9) containing 500 mM NaCl and 1 mM PMSF. After breaking cells by microfluidizer (AH-TITAN PLUS) at 800 bar for 15 min, the lysate was centrifuged at 12,000 rpm for 40 min at 4 °C. The supernatant was transferred to a fresh tube containing 4 mL of 50 % slurry of Streptactin beads 4FF (Smart-Lifesciences). After rotating at 4 °C for 2 h, the mixture was transferred to a column to allow the settlement of the beads. The column was washed with 50 mL lysis buffer, and eluted with 10 mL elution buffer (lysis buffer with 2.5 mM dethiolated biotin). Eluates were concentrated to 50 μL with Amicon Ultra 30K device (Millipore) at 4 °C. Dethiolated biotin was removed through buffer exchange during the ultrafiltration process. Bradford assay was used to measure the final concentration of proteins. The proteins were divided into 5 μL aliquots, snap-frozen in liquid nitrogen and stored at -80 °C.

### In vitro phase separation assays

MBP-3C-GFP-RAT-1 proteins were diluted to desired molarity with different concentrations of NaCl using dilution buffer (20 mM HEPES, 5 % glycerol, pH 8.2) containing various NaCl concentrations. To eliminate MBP, protein solutions were incubated with 1 x HRV 3C protease (Takara) at RT for 10 min. For 1,6-hexanediol assays, MBP-GFP-RAT-1 was diluted to a final concentration of 3 μM in reaction buffer (20 mM HEPES, 5 % PEG-8,000, 150 mM NaCl, 5 % glycerol, pH 8.2) in the presence or absence of 5 % 1,6-hexanediol (Macklin). The solutions were incubated at room temperature for 10 min and then analysed. For phosphorylation assays, purified StrepII-tagged *C. elegans* DYF-5 (human MAK homolog) (0.5 μM), or heat-inactivated DYF-5 (0.5 μM) (after 10-min treatment at 95 ℃), was incubated with MBP-GFP-RAT-1 (2.5 μM) in reaction buffer (50 mM Tris-HCl, 150 mM NaCl, 10 mM MgCl2, 2 mM ATP, pH 8.0) and 1 % PEG-8,000. The mixture was incubated at RT for 5 min before imaging. For RAT-1 and EB3 co-phase separation assays, 10 μM EB3-mCherry and 1 μM MBP-GFP-RAT-1 were incubated together in BRB80 buffer (pH 6.9) containing 150 mM NaCl and 3 % PEG-8,000. The solutions were incubated at room temperature (or 4 / 20 / 37 ℃) for 10 min and analysed by an Axio Observer Z1 microscope (Carl Zeiss).

### Pull-down assay

To examine the interaction between RAT-1 and EB proteins, MBP-GFP-RAT-1-His (WT), MBP-GFP-RAT-1-His (△429-444), and MBP-GFP-His (Control) were cloned into pET21 vectors and expressed in 100-mL culture of BL21 (DE3) bacteria. For each sample, bacterial cells (pellets) were lysed with 0.5-mL cell lysis buffer (Beyotime, #P0013) containing 1 mg/mL lysozyme and 1 mM PMSF, followed by gentle addition of 10 volumes of HEPES buffer (20 mM HEPES, 500 mM NaCl, 5 % glycerol, pH 8.0) and centrifugation at 12,000 rpm for 10 min. The supernatant was transferred to a new tube, as the total lysate. For pull-down assays, 20 μL total lysate was incubated with 1 μM EB3-mCherry-StrepII and 20 μL Streptactin agarose beads (SmartLifesciences) in 200 μL BRB80 buffer at room temperature for 30 min. Following incubation, the beads were pelleted by centrifugation at 1,000 g for 1 min, and were washed three times with 1 mL BRB80 buffer and eluted with BRB80 buffer containing 2.5 mM D-biotin. Eluates or total lysates were analysed by SDS-PAGE followed by Western blotting.

### Disordered score calculation

RAT-1 sequence was obtained from Wormbase (https://wormbase.org/). Disordered score along the RAT-1 sequence was predicted by PONDR (Predictor of Natural Disordered Regions) (https://www.pondr.com/) using VSL2 and VL3-BA predictors.

### Mass spectrometry analysis to identify phosphorylation sites

For in vivo experiments involving HEK293T cells, two 100-mm cell-culture plates of cells (about 107 cells) were employed. For each plate, flag-EGFP-RAT-1 plasmids (10 μg) were subjected for transfection using LipofectamineTM 3000 Transfection Reagent (Invitrogen). After 24 hours, the cells were harvested and lysed in lysis buffer (Beyotime, #P0013) with 1% Triton X-100 and 1mM PMSF. Four volumes of 1xPBS buffer with 5% glycerol, 500 mM NaCl, and 1mM PMSF were gently mixed with the lysates, followed by centrifugation for 10 min at 12,000 rpm at 4 °C. After centrifugation, lysate supernatants were mixed with pre-washed anti-DYKDDDDK (flag) agarose beads (SmartLifesciences; 50 μL bed volume per sample), and incubated at 4 °C on a shaker for 2 h. After incubation, the beads were washed with 1 mL 1xPBS buffer for 3 times, and then eluted with 250 μL 1xPBS buffer containing 0.2 mg/mL 3xFlag peptides (Beyotime). Final eluates were concentrated to 100 μL with Amicon Ultra 30K device (Millipore) at 4 °C, followed by digestion with sequencing grade modified trypsin at 37 ℃ overnight.

For in vitro experiments, purified recombinant MBP-GFP-RAT-1 (20 μM) was incubated with purified *C. elegans* DYF-5 (1 μM) at 30 ℃ for 60 min, in the presence of 50 mM Tris-HCl (pH 8.0), 10 mM MgCl2, 150 mM NaCl, and 2 mM ATP. The phosphorylated samples (5 μL) were then subjected to SDS-PAGE analysis. The targeted gel band (around 130 kD) was excised from the gel, reduced with 5 mM of dithiothreitol (DTT) and alkylated with 11 mM iodoacetamide (IAM), followed by in-gel digestion with sequencing grade modified trypsin at 37 ℃ overnight. The peptides were extracted twice with 0.1% trifluoroacetic acid in 50% acetonitrile aqueous solution for 30 min. The peptide extracts were then centrifuged in a SpeedVac to reduce the volume. Peptides were redissolved in 20 μl 0.1% trifluoroacetic acid and 1 μl of extracted peptides were analyzed by Thermo Scientific Orbitrap Exploris 480. The digestion products were separated by a 95-min gradient elution at a flow rate 0.300 µL/min with a Thermo-Dionex Ultimate 3000 HPLC system which was directly interfaced with the Thermo Scientific Orbitrap Exploris 480 mass spectrometer. The analytical column was a home-made fused silica capillary column (75 µm ID, 350 mm length) packed with C-18 resin (120 Å, 1.9 µm, Dr.Maisch, Ammerbuch, Germany). Mobile phase A consisted of 0.1% formic acid, and mobile phase B consisted of 80% acetonitrile and 0.1% formic acid. The Orbitrap Exploris 480 mass spectrometer was operated in the data-dependent acquisition mode using Xcalibur 4.5 software and there is a single full-scan mass spectrum in the Orbitrap (350-1500 m/z, 60,000 resolution) followed by 20 MS/MS (mass spectrometry / mass spectrometry) scans in the Orbitrap. The MS/MS spectra from each LC-MS/MS run were searched against the RAT-1 protein sequence using Proteome Discoverer (Version PD 3.0, Thermo-Fisher Scientific, USA). The search criteria were set as follows: full tryptic specificity was required; two missed cleavage sites were allowed; Oxidation (M) and phosphorylation (P) were set as variable modification; Carbamidomethyl (C) was set as fixed modification; precursor ion mass tolerances were set at 20 ppm for all MS acquired in an Orbitrap mass analyzer; and the fragment ion mass tolerance was set at 0.02 Da for all MS2 spectra acquired.

### Western blot

For interaction between RAT-1 and EB3, first antibodies were used at the following dilutions: anti-6xHis (Proteintech, #66005-1-Ig), 1:1,000; anti-Strep (Solarbio, #K200012M), 1:5,000.

For DYF-5-mediated phosphorylation assays, purified MBP-GFP-RAT-1 (20 μM) was incubated with *C. elegans* DYF-5 (1 μM) (active or heat-inactivated) at 30 ℃ for 60 min, in the presence of 50 mM Tris-HCl (pH 8.0), 10 mM MgCl2, 150 mM NaCl, and 2 mM ATP. The reaction mixtures were analysed by SDS-PAGE followed by Western blotting. First antibodies included anti-His (Proteintech, #66005-1-Ig; 1:10,000), anti-phosphoserine/threonine (pST) (BD Biosciences, #612548; 1:2,500). After incubation with first antibodies overnight at 4 ℃ on a shaker, the PVDF membranes were washed three times with 1xTBST buffer (10 min per wash on a shaker). Secondary antibodies were added at dilutions of 1:10,000 (goat anti-mouse/rabbit, EasyBio) and incubated for 1 h at room temperature on a shaker, followed by 3 washes with 1xTBST buffer. Finally, chemiluminescence (SuperSignal™ West Pico PLUS, Thermo Fisher Scientific) agent was added and signals were visualized using a FluorChem illuminator (ProteinSimple).

### AlphaFold3-based structural prediction

Protein sequences were obtained from the Uniprot database (https://www.uniprot.org/). The monomer structure of RAT-1, multimer structures of RAT-1 and *Ce*EBP-2 (*C. elegans* EB homolog), RAT-1 and *Hs*EB1 (*H. sapiens* EB1), RAT-1 and *Hs*EB2 (*H. sapiens* EB2), RAT-1 and *Hs*EB3 (*H. sapiens* EB3), were predicted by AlphaFold3 (alphafoldserver.com/) (Abramson et al., 2024). Predicted alignment error (PAE) plots were directly obtained from the AlphaFold3 server. Structural representations were generated using UCSF ChimeraX (https://www.rbvi.ucsf.edu/chimerax/).

### Forward genetic screen and suppressor cloning

We used forward genetic screens to isolate *tba-5 (A19V) (qj14)* suppressors. The mutant animals (P0) were synchronized at the late L4 larval stage, collected in 4 mL M9 buffer, and incubated with 50 mM ethyl methanesulfonate (EMS) for 4 hours at room temperature (RT) with rotation. Animals were then washed with M9 three times and cultured under standard conditions. After 24 hours, animals were bleached. Eggs (F1) were distributed and raised on NGM plates, each containing about 20 eggs. Adult F2 animals were collected and subjected to dye filling. We used sibling subtraction method to clone suppressor mutations (Joseph et al., 2018). Briefly, homozygous suppressor strains were crossed with original *tba-5 (A19V)* mutants, and 10 F2 progenies with (S) or without (NS) suppressor mutations were isolated according to dye-filling phenotype, individually. The progenies (F3) of all S groups were mixed together for whole-genome sequencing (WGS), as well as all NS groups. We compared differential homozygous DNA variants between S and NS groups, and identified suppressor mutations by rescue experiments.

### In vitro microtubule (MT) binding and dynamics assay

For in vitro MT binding assay, GMPCPP-stabilized MT polymerization followed the protocol described previously (Chen et al., 2025). Briefly, MRB80 buffer (BRB80 buffer with 4 mM MgCl2) containing 0.3 μM rhodamine-conjugated tubulins (Cytoskeleton), 5.7 μM unlabeled porcine brain tubulins (Cytoskeleton), and 1 mM GMPCPP (JenaBiosciences) were incubated at 37 ℃ for 30 min. After centrifugation at 15,000 rpm for 30 min, the pellet (containing polymerized MTs) was suspended with MRB80 buffer, and then flowed into anti-β-tubulin enveloped chambers. Next, 10 μM MBP-GFP-RAT-1 in the presence of 150 mM NaCl was flowed into the chambers and incubated for 5 min, followed by extensive wash by HEPES buffer (20 mM HEPES, 150 mM NaCl, pH 8.2). The samples were analysed by an Axio Observer Z1 microscope (Carl Zeiss).

In vitro MT dynamics assay followed the protocol described previously (Zhang et al., 2022). In brief, the flow chamber was incubated first with 0.2 mg/mL PLL-PEG-biotin (Susos AG) and then with 1 mg/mL neutravidin (Invitrogen) in MRB80 buffer. Subsequently, rhodamine-conjugated and biotin-conjugated MT seeds, or HiLyte647-conjugated and biotin-conjugated MT seeds, were attached to the coverslip through biotin–neutravidin interactions. The flow chamber was further blocked with 1 mg/mL κ-casein. The reaction mixture, which consisted of MRB80 buffer containing MBP-GFP-RAT-1 (100 nM or 20 μM, optional), EB3-mCherry (50 nM, optional), porcine brain tubulins (10 μM) (Cytoskeleton), 2 mM GTP, 0.2 mg/ml κ-casein, 1% PEG-8,000 and oxygen scavenger mix (50 mM glucose, 400 μg/ml glucose oxidase, 200 μg/ml catalase, and 4 mM DTT) was added to the chamber. For fluorescent tubulins, the percentage of HiLyte488-conjugated tubulins is 12.5%, and the percentage of rhodamine-conjugated tubulins (Cytoskeleton, Inc) is 2.5%. The flow chamber was sealed with vacuum grease and imaged immediately at 30 °C using a Nikon Eclipse Ti-E inverted microscopy equipped with a 100×/1.4 objective. Images were acquired at the TIRF (total internal reflection fluorescence) mode with reflection angle of 65°, laser wavelength of 488 nm / 561 nm / 640 nm and 8 sec intervals.

### Quantification and statistical analysis

The sample sizes were determined based on prior published studies of similar design. GraphPad Prism 9 (GraphPad Software, Inc.) was used for statistical analyses. For comparisons between two groups exhibiting Gaussian distribution, two-sided unpaired t tests were applied. For non-normally distributed data, two-sided Mann-Whitney U tests were employed. For comparisons between more than 2 groups with Gaussian distribution and equivalent standard deviation, two-sided one-way ANOVA with Dunnett’s (when comparing with one control group) or Tukey’s (when comparing with each other) multiple comparisons tests were applied. For multiple groups with Gaussian distribution but inequivalent standard deviation, two-sided Welch’s ANOVA with Dunnett’s or Tukey’s tests were performed. For multiple groups without Gaussian distribution, two-sided nonparametric tests (Kruskal-Wallis test with Dunn’s multiple comparisons) were applied. N represents the number of samples per group. Data are presented as mean ± SD unless stated otherwise. Statistical significances are designated as: ns *P* > 0.05, * *P* < 0.05, ** *P* < 0.01, *** *P* < 0.001 and **** *P* < 0.0001.

## Acknowledgements

This work was supported by the National Key R&D Program of China (2024YFA1307301, 2022YFA1302700) and the National Natural Science Foundation of China (92254306 to G.O.; 323B200173 to K.X.; 32270773 and 32470730 to W.L.; 32430026, 32021002, and 31991191 to G.O.). We thank Dr. Kai Jiang (Wuhan University) and Dr. Xin Liang (THU) for their assistance with microtubule dynamics assays. We thank Dr. Pilong Li (THU), Dr. Xuebiao Yao (USTC), Dr. Shengqi Xiang (USTC), Dr. Mingjie Zhang (HKUST) and Dr. Gaofeng Pei (THU) for their insightful discussions on phase separation. We thank Dr. Andrew Carter (MRC-LMB) and Dr. Carsten Janke (Institut Curie) for their insightful discussions on cilia and microtubules. We thank Dr. Shanshan Xie (Zhejiang University) for kindly sharing the hTERT-RPE1 cell line. We thank Wenwan Rong and Dr. Haiteng Deng in Technology Center for Protein Science, Tsinghua University for mass spectrometry analysis. We also thank the Facility of Cell Imaging at the Tsinghua University Technology Center for Protein Research, and Nikon Bioimaging Center (especially Jinyu Wang) for their technical assistance with imaging.

## Author Contributions

G.O. and K.X. conceived and designed this study. G.O. supervised this study. K.X. performed all experiments. Z.Z. provided *Ce*DYF-5 proteins. K.X. analysed the data. K.X., G.O., W.L. and Y.C. wrote this manuscript.

## Data Availability

Data used in the study are included in the manuscript and/or supporting information.

**Movie S1 (separate file). Time-lapse live-cell imaging showing intra-ciliary dynamics of GFP::RAT-1 and DYF-11::mScarlet.** This movie corresponds to Fig. S2B. Total time: 1 min (3.75× sped up).

**Movie S2 (separate file). Time-lapse live-cell imaging showing fusion (left) and fission (right) events of GFP::RAT-1 condensates within cilia.** This movie corresponds to Figs. 2G-H. Total time: 6 sec.

**Movie S3 (separate file). Time-lapse TIRF imaging of in vitro microtubule dynamics with RAT-1 and EB3.** This includes GMPCPP-stabilized seed MTs (indicated by cyan arrowheads), porcine brain tubulins (10 μM, HiLyte488-conjugated or rhodamine-conjugated), EB3-mCherry (50 nM), and MBP-GFP-RAT-1 (100 nM). This movie corresponds to Fig. 6H. Total time: 10 min (120× sped up).

**Dataset S1 (separate file).** All the *C. elegans* strains used in this study.

---

**Fig. S1.**
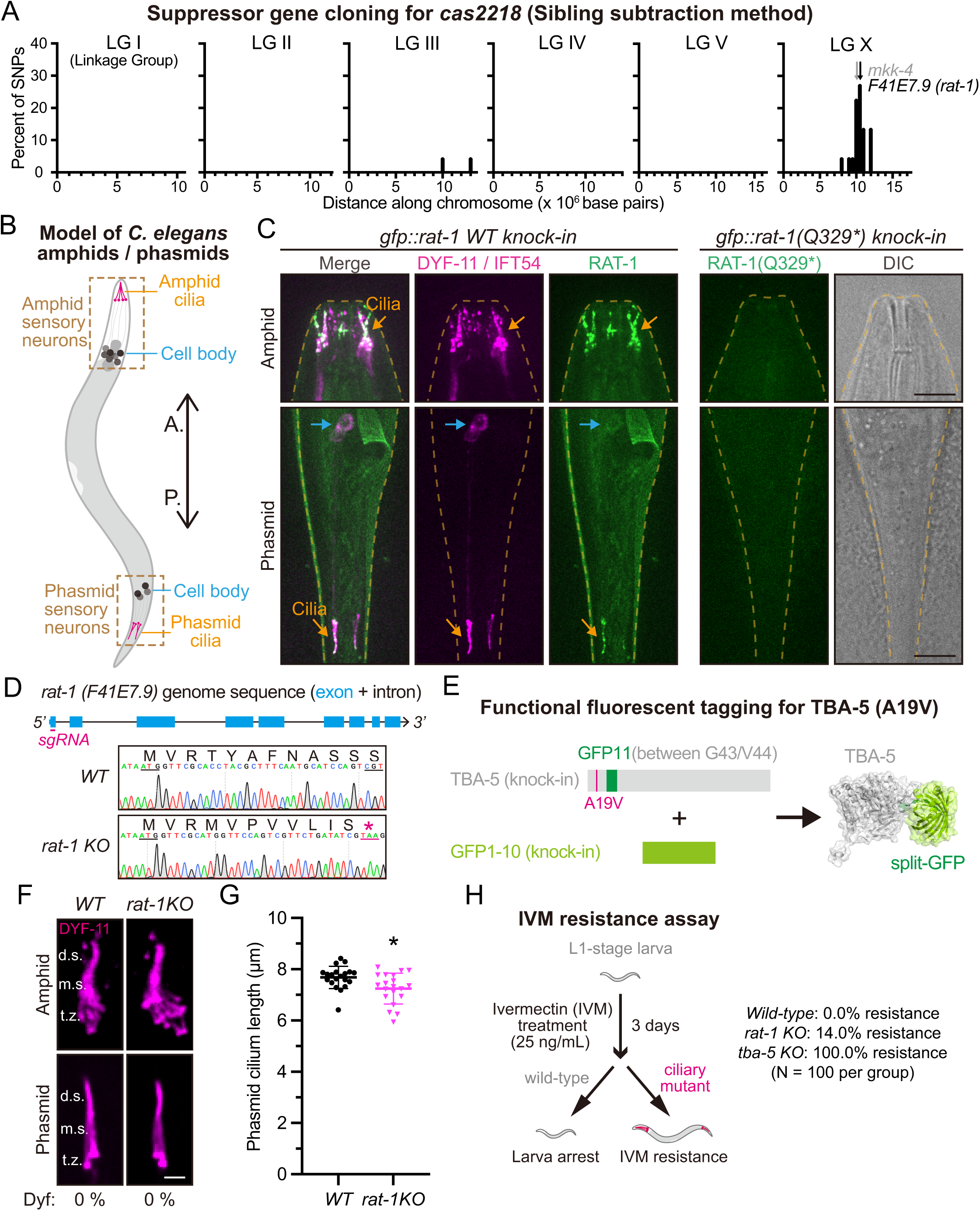
RAT-1 (F41E7.9) is a ciliary protein essential for cilia integrity. (A) Percentage of candidate single nucleotide polymorphisms (SNPs) along the whole genome (linkage group I, II, III, IV, V and X) of *C. elegans* (*cas2218*) using the sibling subtraction method, a methodology for genetic suppressor cloning (Joseph et al., 2018). In the linkage group (or chromosome) X, only two SNPs locating in *mkk-4* and *F41E7.9* (*rat-1*), respectively, belong to change-of-function mutations (missense mutation or premature stop codon). (B) Schematic of *C. elegans* amphid and phasmid sensory neurons and cilia. A., anterior; P., posterior. (C) Representative images of amphid and phasmid cilia in wild-type *gfp::rat-1* (left) or *gfp::rat-1 (Q329*) (cas2218)* knock-in animals. Cilia are marked with DYF-11(IFT54)::mScarlet. Cell bodies are indicated in blue, and cilia in orange. Scale bar, 10 μm. (D) Schematic showing generation of the *rat-1* knockout (KO) allele by CRISPR-Cas9-mediated genome editing. sgRNA targets the first exon of *rat-1* genome sequence. Sanger sequencing confirmed the generation of homozygous *rat-1* KO strains. (E) Schematic of the split-GFP-based functional tagging strategy to label endogenous TBA-5 (WT or the A19V mutant) (Xu et al., 2024). (F) Representative images of phasmid cilia in wild-type or *rat-1* KO worms. Cilia are marked with DYF-11(IFT54)::mScarlet. Percentage of dye-filling defective (Dyf) animals is indicated below the images. t.z., transition zone; m.s., middle segment; d.s., distal segment. Scale bar, 2 μm. (G) Quantification of phasmid cilia length in each group in (F). N = 20 phasmids. Shown as mean ± SD. Unpaired t tests were performed. * *P* < 0.05. (H) Ivermectin (IVM) resistance assay showing that 14 % *rat-1* KO animals exhibited IVM resistance, a representative phenotype of ciliary mutants.

**Fig. S2.**
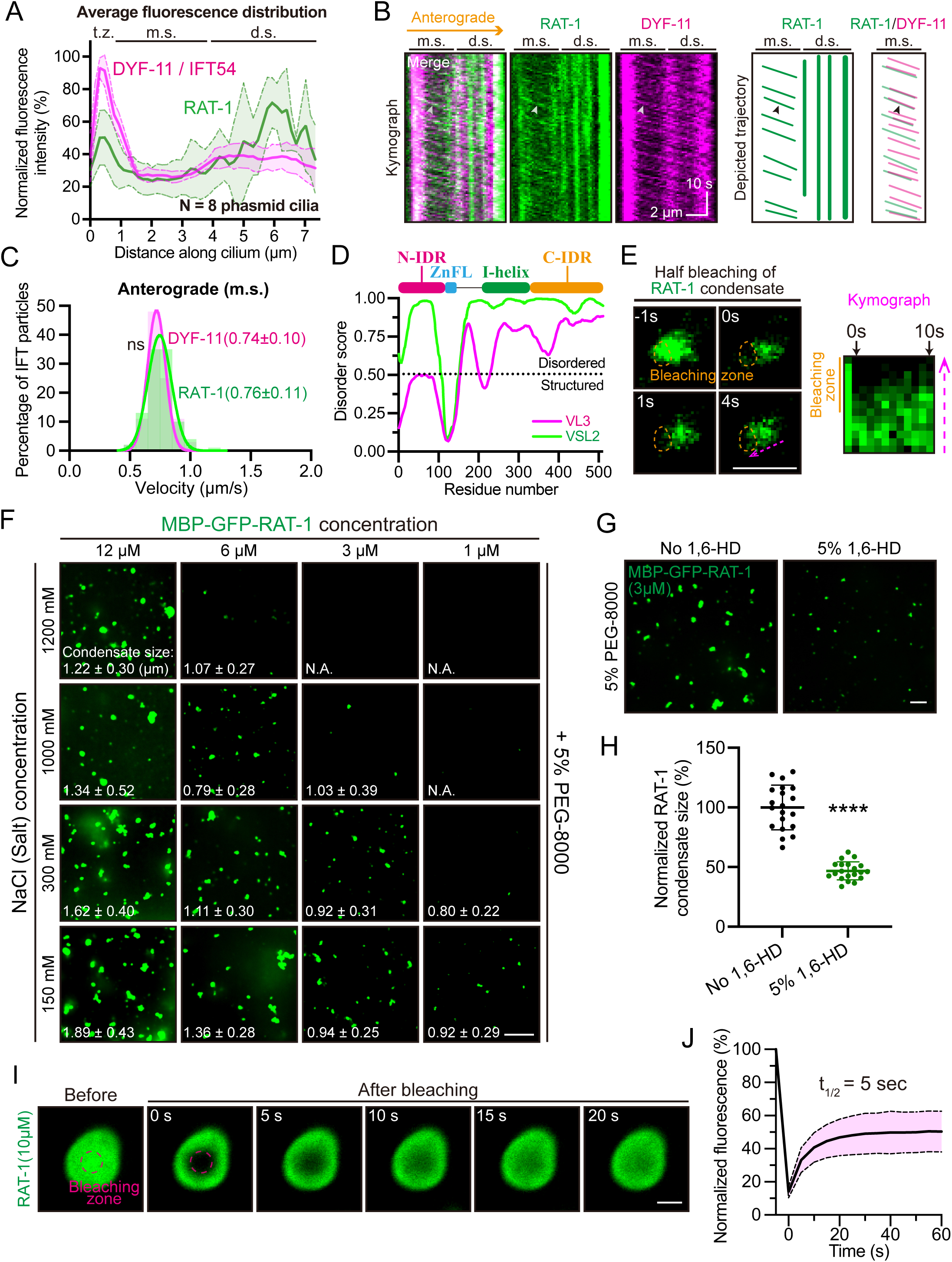
RAT-1 undergoes intraflagellar transport (IFT) and forms condensates in cilia. (A) Distribution of RAT-1 and DYF-11 fluorescence along the phasmid cilia in *gfp::rat-1; dyf-11::mScarlet* animals. 8 phasmid cilia were used for quantification and the average fluorescence was shown. The maximal fluorescence in each cilia was normalized to 100%. Shown as mean ± SD. t.z., transition zone; m.s., middle segment; d.s., distal segment. (B) (Left) Representative kymographs showing concurrent movements of RAT-1 and DYF-11 within phasmid cilia. White arrowhead indicates a representative IFT trajectory with co-localization of RAT-1 and DYF-11. (Right) Corresponding trajectories extracted from the kymograph, showing movements of RAT-1 or co-migrating RAT-1 and DYF-11. m.s., middle segment; d.s., distal segment. Scale bar, 2 μm (horizontal) and 10 s (vertical). See also Movie S1. (C) Histogram showing distribution of non-condensate RAT-1 or IFT (represented by DYF-11) velocities within ciliary middle segments (m.s.). Data were shown as mean ± SD with Gaussian curves. Unpaired t tests were performed to compare the differences. N = 100 per group. (D) Predicted disordered score along the primary sequence of RAT-1 by PONDR using VSL2 and VL3-BA predictors. (E) (Left) Representative FRAP (fluorescence recovery after photobleaching) images of GFP::RAT-1 condensates within cilia. A half condensate was bleached (indicated in orange). Scale bar, 1 μm. (Right) Kymograph showing rapid FRAP dynamics of the RAT-1 condensate. Plot was generated along the magenta arrow. (F) Phase diagram showing the condensate formation capacity of RAT-1 depends on both protein and salt concentrations. Quantification of condensate size was shown as mean ± SD (N = 20 condensates from 3 independent experiments). N.A., not applicable. Scale bar, 5 μm. (G) Representative images of in vitro condensate formation of RAT-1 with or without 5 % 1,6-hexanediol (1,6-HD), in the presence of 150 mM NaCl. Scale bar, 5 μm. (H) Quantification of RAT-1 condensate size with or without 5 % 1,6-hexanediol (1,6-HD). N = 20 condensates from 3 independent experiments. Unpaired t tests were performed. **** *P* < 0.0001. (I) Representative FRAP images of GFP-RAT-1 condensates in vitro. MBP tag was removed. Bleaching zone was indicated in magenta. Scale bar, 1 μm. (J) Normalized fluorescence recovery of RAT-1 condensates in vitro after photobleaching. Shown as mean ± 95% CI (confidence interval). N = 10.

**Fig. S3.**
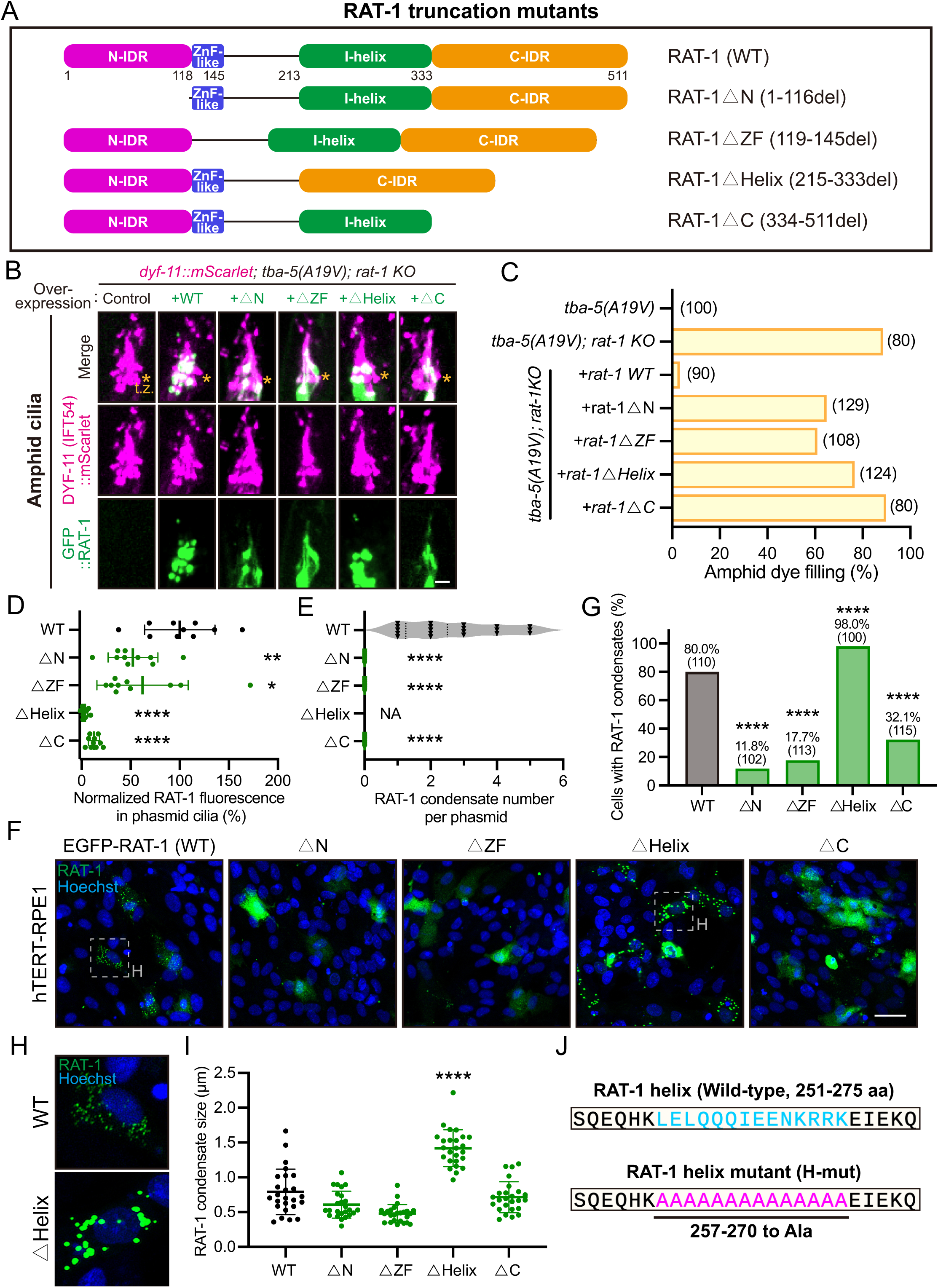
The internal helix (I-helix) of RAT-1 auto-inhibits its condensation formation capacity. (A) Schematic of RAT-1 truncation mutants. (B) Representative images of amphid cilia in *tba-5 (A19V); rat-1 KO* animals overexpressing wild-type GFP::RAT-1 or truncation mutants. Cilia are marked with DYF-11(IFT54)::mScarlet (knock-in). t.z., transition zone (indicated by orange asterisks). Scale bar, 2 μm. (C) Percentage of amphid dye-filling positive animals in each group in (B). (D) Quantification of total GFP::RAT-1 fluorescence within phasmid cilia. The mean value of WT group is normalized to 100 %. N = 10 phasmids. Shown as mean ± SD. Mann-Whitney tests (Nonparametric statistics) were performed. * *P* < 0.05; ** *P* < 0.01; **** *P* < 0.0001. (E) Quantification of GFP::RAT-1 condensate numbers within phasmids. Mann-Whitney tests (Nonparametric statistics) were performed. NA, not applicable; **** *P* < 0.0001. (F) Representative images showing overexpressed EGFP-RAT-1 truncation mutants in hTERT-RPE1 cells. Scale bar, 10 μm. (G) Percentage of EGFP-RAT-1 transfection-positive cells forming liquid-like condensates in each group in (F). Chi-square tests were performed. **** *P* < 0.0001. (H) Enlarged images in WT or △Helix group in (F). (I) Quantification of EGFP-RAT-1 condensate sizes in each group in (F). N = 25 condensates from 3 independent experiments. Shown as mean ± SD. Unpaired t tests were performed. **** *P* < 0.0001. (J) Local RAT-1 protein sequence (251-275 aa) within the internal helix (I-helix). All amino acids from L257 to K270 were mutated to alanine (A) in the H-mut.

**Fig. S4.**
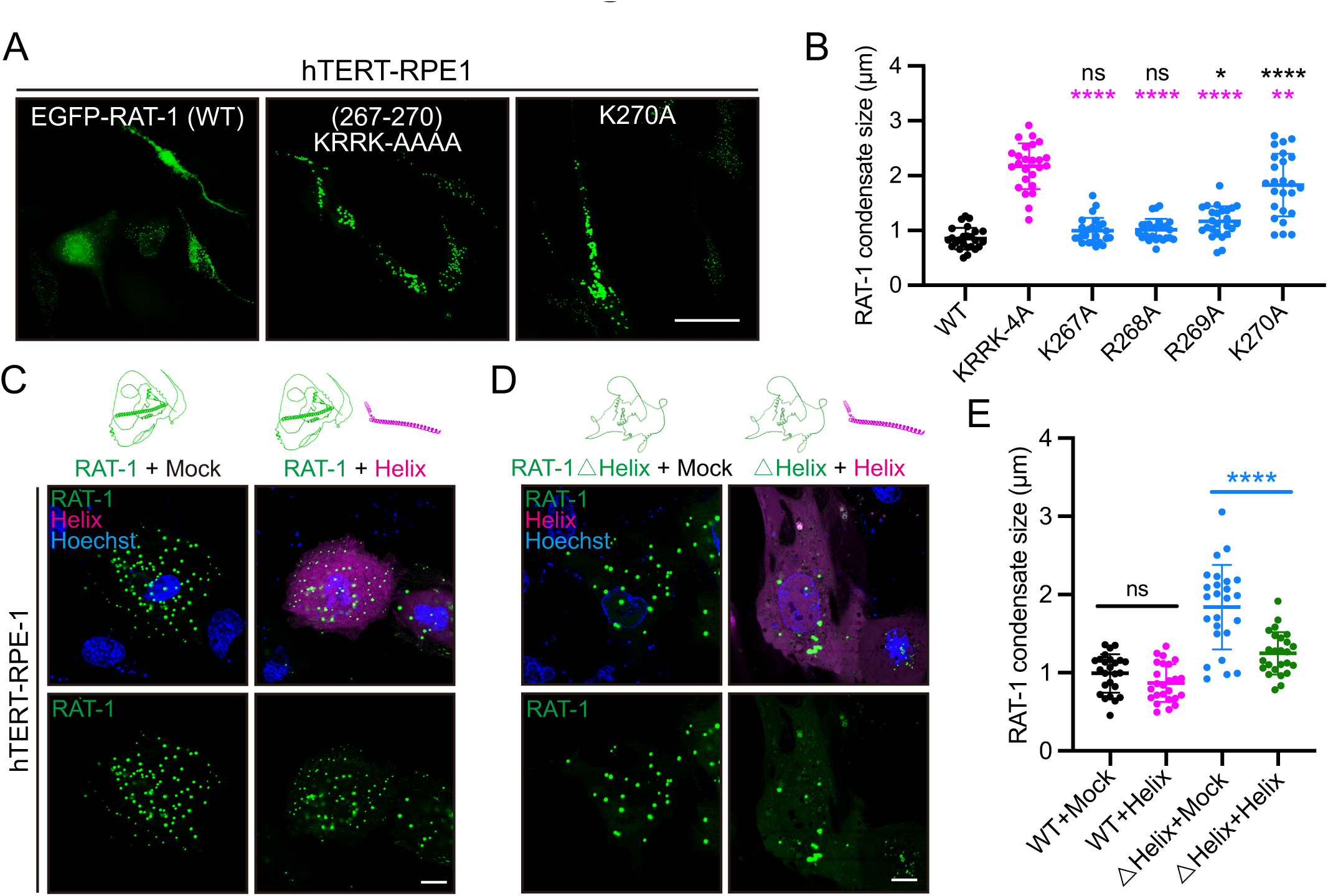
The RAT-1 I-helix inhibits condensate formation of the RAT-1 (△Helix) mutant. (A) Representative live-cell images showing overexpressed WT EGFP-RAT-1, or KRRK-AAAA mutant, or K270A mutant, in hTERT-RPE1 cells. Scale bar, 50 μm. (B) Quantification of EGFP::RAT-1 (WT or mutants) condensate sizes in each group. N = 25 condensates from 3 independent experiments. Shown as mean ± SD. One-way ANOVA with Tukey’s tests were performed. ns, not significant; * *P* < 0.05; ** *P* < 0.01; **** *P* < 0.0001. (C-D) Representative live-cell images showing overexpressed WT EGFP-RAT-1 (C) or the △Helix mutant (D), in the absence (mock) or presence of mCherry-helix (magenta). Scale bar, 10 μm. (E) Quantification of RAT-1 condensate sizes in each group in (C-D). N = 25 condensates from 3 independent experiments. Shown as mean ± SD. Unpaired t tests were performed. ns, not significant; **** *P* < 0.0001.

**Fig. S5.**
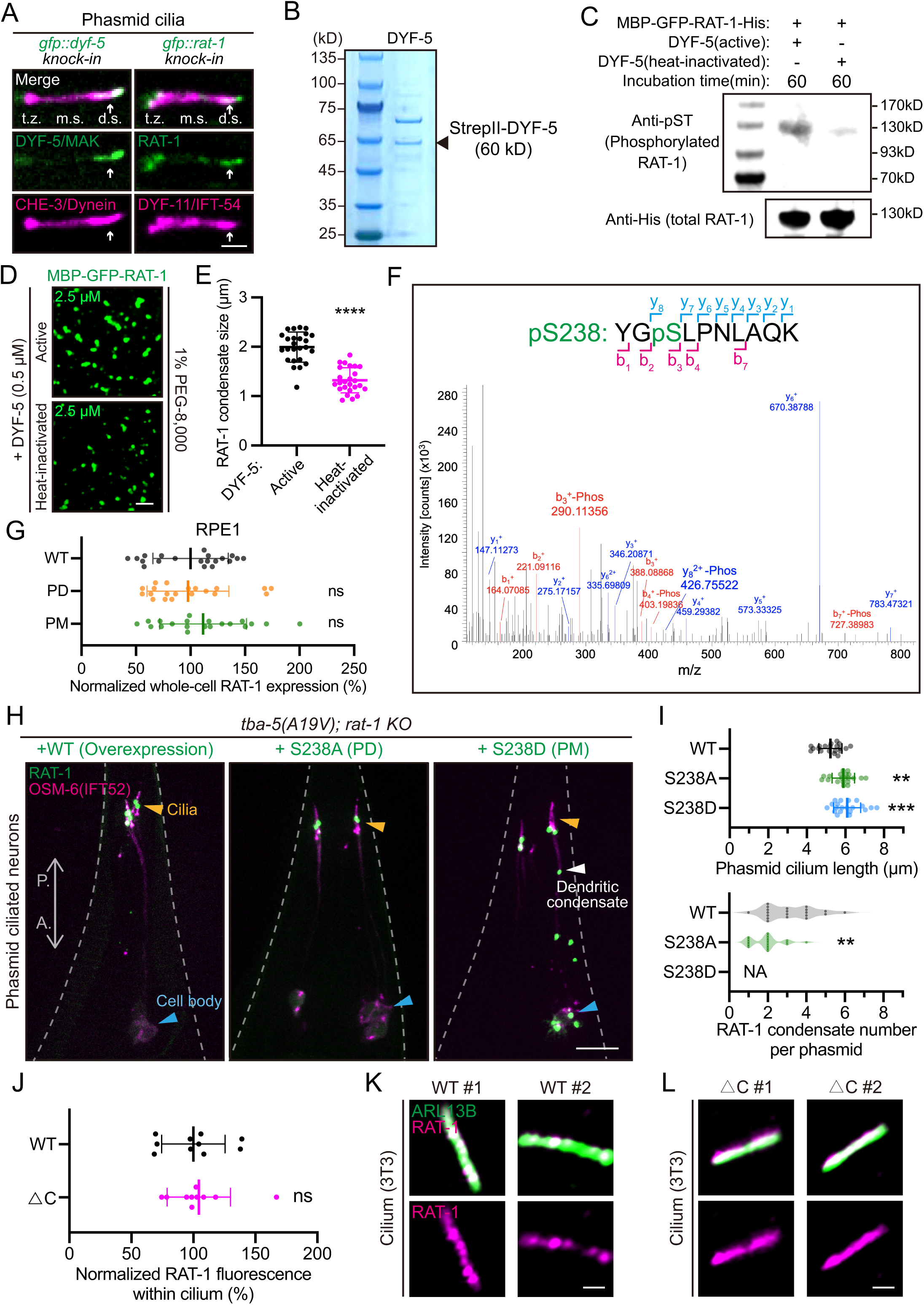
Phosphorylation of the RAT-1 I-helix promotes condensate formation. (A) Representative images of *C. elegans* phasmid cilia in *gfp::dyf-5* knock-in (left) or *gfp::rat-1* knock-in (right) animals. CHE-3(dynein heavy chain)::mScarlet (knock-in) or DYF-11(IFT54)::mScarlet (knock-in) marks the cilia, respectively. t.z., transition zone; m.s., middle segment; d.s., distal segment (indicated by white arrows). Scale bar, 2 μm. (B) SDS-PAGE analysis of recombinant StrepII-DYF-5 purification. The expected molecular weight of full-length StrepII-DYF-5 is about 60 kDa. An additional co-purifying band is observed, the identity of which is currently unknown. The major band corresponding to StrepII-DYF-5 is indicated. (C) Western blot analysis showing in vitro phosphorylation (anti-phosphoserine/threonine) of RAT-1 (131 kD, MBP-GFP-RAT-1-His) by *C. elegans* (Ce) DYF-5/MAK (0.5 μM). Total RAT-1 (anti-His) served as loading control. (D) Representative images of in vitro condensate formation of RAT-1 treated by active CeDYF-5 (top, 0.5 μM), or heat-inactivated CeDYF-5 (bottom, 0.5 μM). Scale bar, 5 μm. (E) Quantification of RAT-1 condensate sizes in each group in (D). N = 25 condensates from 3 independent experiments. Shown as mean ± SD. Unpaired t tests were performed. **** *P* < 0.0001. (F) LC-MS/MS (mass spectrometry) result of the determination of RAT-1 phosphorylation site (S238). (G) Quantification of whole-cell RAT-1 overexpression levels in each group in Fig. 3J. The mean value of WT group is normalized to 100 %. N = 20 cells from 3 independent experiments. ns, not significant. (H) Representative images of phasmid ciliated neurons in *tba-5 (A19V); rat-1 KO* animals overexpressing WT GFP::RAT-1, or the S238A (PD), or the S238D (PM) mutant. Cell bodies of the ciliated neurons are indicated in blue, dendritic RAT-1 condensates in white, and cilia in orange. A., anterior; P., posterior. Scale bar, 10 μm. (I) Quantification of phasmid cilia length (top, one-way ANOVA with Dunnett’s tests) or RAT-1 condensate number within phasmid cilia (bottom, Mann-Whitney tests) in each group. N = 20 phasmids per group. NA, not applicable. (J) Quantification of total RAT-1 fluorescence within the cilia of NIH-3T3 cells. The average value of WT group is normalized to 100 %. N = 10 cilia. Shown as mean ± SD. Unpaired t tests were performed. ns, not significant; ** *P* < 0.01; *** *P* < 0.001. (K-L) Representative immunofluorescence images showing localization of WT Flag-RAT-1 (K) or the △C mutant (L) within the cilia of two independent NIH-3T3 cells (#1 and #2). Scale bar, 1 μm.

**Fig. S6.**
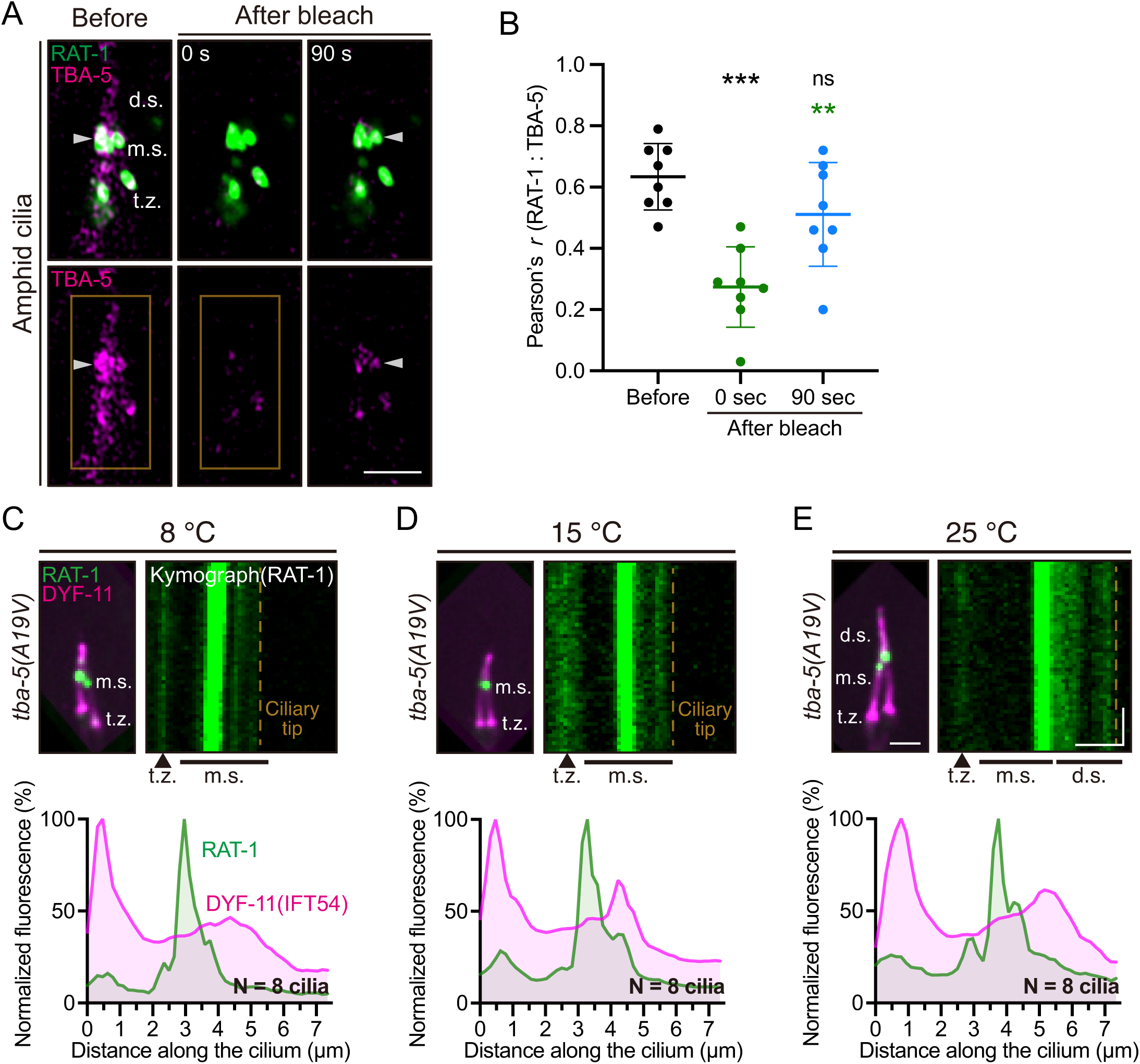
RAT-1 condensates enrich α-tubulin TBA-5 in cilia. (A) Representative FRAP (fluorescence recovery after photobleaching) images of GFP::RAT-1 and TBA-5::mScarlet (overexpression) within the amphid cilia of wild-type animals. White arrowheads indicate the RAT-1 condensates in cilia. Bleaching region is marked in orange. t.z., transition zone; m.s., middle segment; d.s., distal segment. Scale bar, 2 μm. (B) Pearson’s *r* between RAT-1 and TBA-5 fluorescence intensities in amphid cilia. N = 8 cilia per group. Shown as mean ± SD. One-way ANOVA with Tukey’s tests were performed. ns, not significant; ** *P* < 0.01; *** *P* < 0.001. (C-E) (Top) Representative live image (left) and kymograph (right) of *C. elegans* phasmid cilia in *tba-5 (A19V); gfp::rat-1* knock-in animals at 8 ℃ (C), 15 ℃ (D) or 25 ℃ (E). Ciliary tips were indicated by orange dashed lines. (Bottom) Distribution of RAT-1 and DYF-11 fluorescence along the phasmid cilia. N = 8 phasmid cilia per group, and the average fluorescence was shown.

**Fig. S7.**
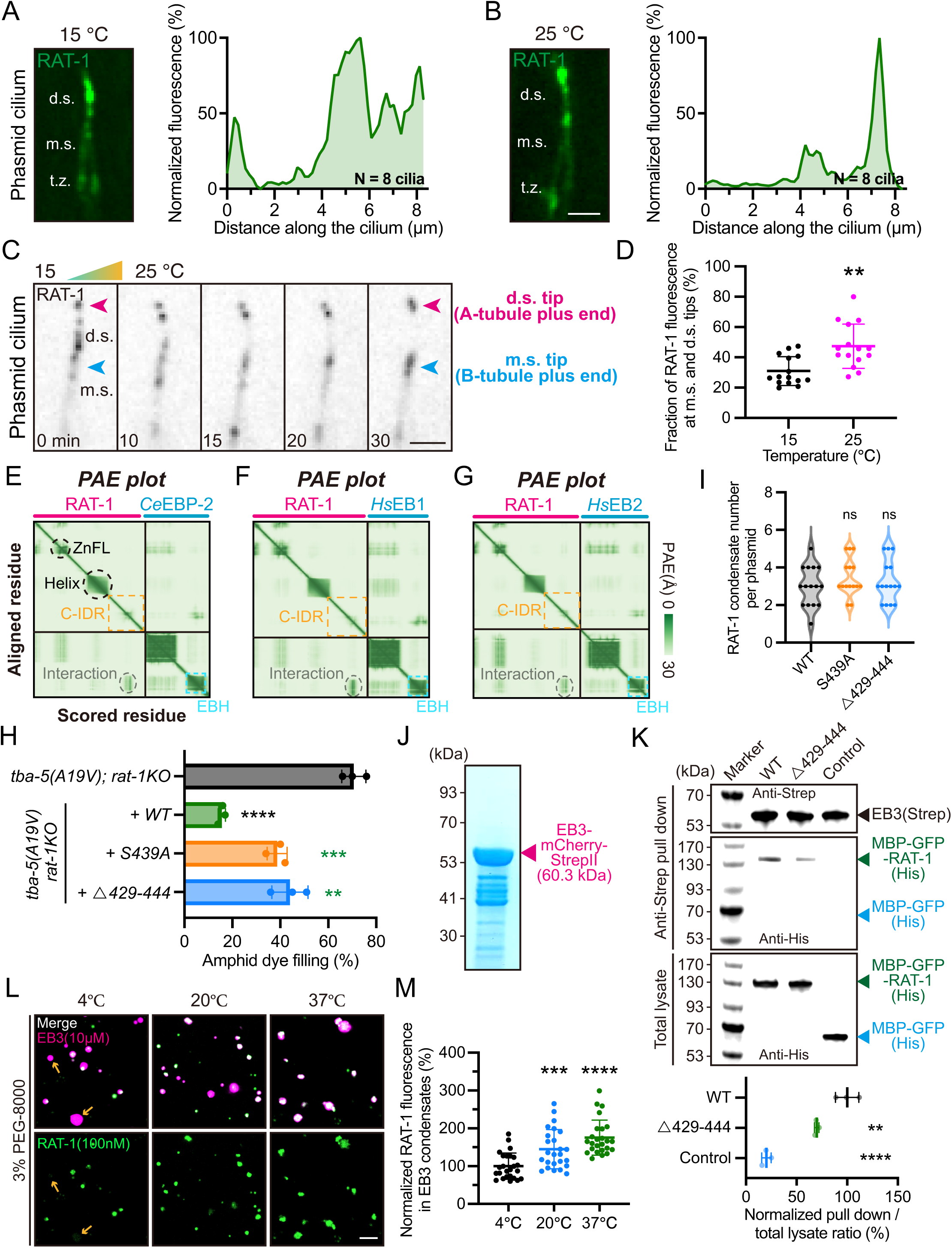
RAT-1 interacts with end-binding (EB) family proteins. (A-B) (Left) Representative live images of *C. elegans* phasmid cilia in *gfp::rat-1* knock-in animals at 15 ℃ (A) or 25 ℃ (B). (Right) Normalized distribution of RAT-1 fluorescence along the cilia. 8 phasmid cilia were used for quantification and the average fluorescence was shown. The maximal fluorescence was normalized to 100%. Scale bar, 2 μm. (C) Time-lapse images showing temperature-dependent dynamics of GFP::RAT-1 within *C. elegans* phasmid cilia. m.s. (middle segment) tip and d.s. (distal segment) tip were indicated by blue and magenta arrowheads, which represent the plus ends of B-and A-tubules of ciliary doublet microtubules, respectively. Scale bar, 2 μm. (D) Quantification of the fraction of RAT-1 fluorescence at m.s. and d.s. tips (assumed microtubule plus ends) along the entire cilia. N = 15 phasmids. Shown as mean ± SD. Unpaired t tests were performed with Welch’s correction. (E-G) AlphaFold3-predicted PAE (predicted alignment error) plots of RAT-1 and *C. elegans* (Ce) EBP-2 (E), RAT-1 and human (Hs) EB1 (F), and RAT-1 and human (Hs) EB2 (G). C-IDR was indicated in orange. EB homology domain (EBH) was indicated in blue. (H) Percentage of amphid dye-filling positive animals at 15 ℃ in each group in Fig. 6D. N = 100 animals per replicate. N = 3 independent replicates. Unpaired t tests were performed. (I) Quantification of RAT-1 condensate numbers in phasmids in each group in Fig. 6D. N = 15 phasmids. Mann-Whitney tests (Nonparametric statistics) were performed. ns, not significant. (J) SDS-PAGE analysis showing purification of EB-mCherry-strepII. (K) (Top) Western blot analyses of EB3-mCherry-StrepII (60.3 kD) and wild-type MBP-GFP-RAT-1-His (130.9 kD), the △429-444 mutant (129.1 kD), or MBP-GFP-His (control, 71.7 kD) following anti-StrepII pull-down assays (see Methods). Equal amounts of RAT-1 or control proteins were used in each assay, as confirmed through anti-His immunoblotting of total lysates. (Bottom) Quantification of wild-type MBP-GFP-RAT-1, the △429-444 mutant, and MBP-GFP control proteins in pull-down samples relative to their corresponding total lysates. Data are presented as the pull-down / total lysate ratio, with the average value of the WT group normalized to 100 %. N = 3 independent replicates. One-way ANOVA with Dunnett’s tests were performed. (L) Representative images of in vitro co-phase separation of MBP-GFP-RAT-1 (100 nM) and EB3-mCherry (10 μM) at different temperatures. Orange arrows indicated EB3 condensates without apparent RAT-1 localization. Scale bar, 5 μm. (M) Quantification of RAT-1 fluorescence in EB3 condensates at different temperatures. The mean value of 4 ℃ group is normalized to 100 %. N = 25 condensates from 3 independent experiments. Shown as mean ± SD. One-way ANOVA with Dunnett’s tests were performed. ** *P* < 0.01; *** *P* < 0.001; **** *P* < 0.0001.

